# Older adults amplify passive gait stability during obstacle crossing without weakening the stabilizing synergy

**DOI:** 10.64898/2026.06.22.733751

**Authors:** Ashwini Kulkarni, Chuyi Cui, Shirley Rietdyk, Satyajit Ambike

**Affiliations:** School of Rehabilitation Sciences, Old Dominion University, Norfolk, USA; George W Woodruff School of Mechanical Engineering, Georgia Institute of Technology, Atlanta, USA; Department of Health and Kinesiology, Purdue University, West Lafayette, USA

**Keywords:** margin of stability, ageing, fall prevention, motor synergy, proactive gait control, locomotion

## Abstract

Older adults sustain disproportionately severe injuries from trip-induced falls during obstacle crossing. Such falls depend partly on forward momentum when the foot crosses the obstacle. MOS_AP_, an index of passive dynamic gait stability, reflects this momentum. We quantified MOS_AP_ and a synergy index from uncontrolled manifold analysis of step length and extrapolated center of mass in 25 young (21.6 ± 3.5 yr) and 23 older adults (68 ± 4.3 yr) during unobstructed and obstructed walking, to test whether MOS_AP_ increases during obstacle crossing and whether it is actively stabilized at each step. Both groups increased MOS_AP_ progressively over two approach steps by reducing forward momentum and shifting the center of mass posteriorly. Older adults showed greater increases at the crossing steps. The synergy index was positive for all steps, showing that deviations in step length and extrapolated center of mass covaried to stabilize MOS_AP_ at step-specific values. The synergy index was not influenced by age. We conclude that adults actively recruit passive body mechanics while approaching and crossing obstacles to reduce the risk of a trip becoming a fall. Older adults amplify this strategy to compensate for diminished neuromuscular corrective capabilities.

## 1. Introduction

Older individuals frequently use external aids to improve their movements and reduce risk of falls and injuries. As age increases, individuals rely more on external support, initially gripping the railing on stairs and pushing against armrests to rise from chairs, and eventually using walkers for mobility. In addition to such overt adaptations there may be covert changes in how older adults modulate and use their body mechanics to stay safe and mobile. Young adults demonstrate such behaviors: while approaching obstacles, they gradually shift their body mechanics over several approach steps, creating a configuration that minimizes the likelihood of tripping as well as fall risk should they trip [1, 2]. Here, we address whether older adults employ the same proactive strategy, perhaps even more extensively given that age-related decline in neuromuscular and sensory-motor capacity for corrective responses [3, 4] makes them more vulnerable to trip-induced falls and subsequent injuries [5, 6].

We address this question by investigating the margin of stability in the anterior-posterior direction (MOS_AP_). MOS_AP_ quantifies the instantaneous passive dynamic stability during locomotion by measuring the relationship between the extrapolated center of mass (XcoM), a projection of the center of mass that accounts for its velocity, and the base of support (BOS) boundary at heel contact [7]. While walking on a clear walkway, both younger and older adults typically maintain MOS_AP_ values indicating that the XcoM is ahead of the BOS, which signals efficient forward progression with low muscular effort [8–10]. This walking pattern requires less energy [11, 12] but provides less stability against forward falls, a liability for locomotor tasks imposing forward perturbations. In such scenarios, a safer approach would be to increase passive stability and accept higher energy costs for those steps.

This is indeed observed in human locomotion in contexts such as obstacle crossing. When stepping over an obstacle, young adults increase passive dynamic stability compared to unobstructed walking [1] and older adults demonstrate higher passive stability than younger adults [13]. Our previous work revealed two additional key observations: young adults adjust MOS_AP_ gradually over two approach steps rather than abruptly at the obstacle, and they stabilize MOS_AP_ at each of those steps through coordinated adjustments of body state and foot placement.

We emphasize a key operational distinction: MOS_AP_ quantifies the passive dynamic stability of gait that emerges solely from the physics of body motion. A systematic shift in that stability ahead of a known hazard, in contrast, signals active neural control. Furthermore, we used a synergy index derived from the uncontrolled manifold method [14] to ask whether MOS_AP_ is regulated around its selected value via covariation between step length and XcoM. By analogy, MOS_AP_ is a speeding bicycle’s natural stability arising from its momentum and structure. Shifting MOS_AP_ is like a rider slowing down before a turn. The covariation our synergy index detects is like the rider’s steering corrections to remain upright.

We propose three hypotheses on how aging affects obstacle navigation. The first hypothesis addresses passive dynamic stability, with the magnitude of MOS_AP_ as the dependent variable, whereas the other two address the stabilization of MOS_AP_ itself, with the synergy index as the dependent variable.

When navigating obstacles, MOS_AP_ will be higher compared to a clear walkway, it will change across two approach steps and the obstacle-crossing steps, with older adults demonstrating greater increase and higher stability, resulting in a walkway × step × age interaction (H1).

Second, based on our observations of robust synergies in young adults and greater caution in older adults, we hypothesize that both groups will stabilize MOS_AP_ through synergistic coordination of XcoM and foot placement for each step on both walkways (H2).

Third, we expect this synergy to weaken during obstacle crossing compared to the clear walkway, with a greater decline in older adults. Obstacle crossing demands greater limb elevation, requiring increased muscle activations that introduce more signal-dependent noise [15]. This mechanical challenge explains why locomotor synergies weaken during more difficult tasks [1]. The effect is magnified by aging: while older adults maintain synergies on uneven surfaces [16], they show weaker synergies during the more demanding task of obstacle crossing [17].

Previous research examining synergies during obstacle crossing has exclusively focused on the crossing steps alone [17, 18], and not the approach phase where adaptive control begins. We examine behavior across both approach and crossing steps to show the evolution of control that previous work has not captured. We hypothesize that the synergy stabilizing MOS_AP_ will weaken during obstacle crossing for both groups, with older adults showing greater synergy decline. These effects will manifest across multiple approach and crossing steps, resulting in a walkway × step × age interaction (H3).

## 2. Methods

### 2.1 Participants

Sixty healthy young and older adults participated in the study. We excluded five young adults and seven older adults due to poor kinematic tracking. Therefore, data from 25 young (21.6 ± 3.5 years, 68.4 ± 13.3 kg, 1.68 ± 0.09 m, 15 women) and 23 older adults (68 ± 4.3 years, 83.6 ± 17.1 kg, 1.69 ± 0.09 m, 14 women) were included. All participants walked without aid, had no orthopedic, neuromuscular, or dementia disorders, and were independent in daily activities. Vision was normal or corrected-to-normal. The study was approved by the University’s Institutional Review Board (Protocol: IRB-2021-331), and data was collected from May 2021 to May 2022. All participants provided written informed consent.

### 2.2 Equipment and Procedures

Participants walked at their self-selected speed along a 6.0 m walkway, either unobstructed or with an obstacle to step over (Fig. 1A). The obstacle, made of black Masonite and designed to tip if contacted, measured 100 cm wide by 0.4 cm deep, with its height set to 25% of each participant’s leg length. We determined a starting position for each participant such that they took five steps before reaching the obstacle, crossed it with their right leg first, and then took three to four additional steps before stopping. Participants completed two sets of 20 walking trials: first without an obstacle (clear walkway) followed by another with an obstacle (obstructed walkway). We used Vicon Vero motion capture system (Oxford, UK) to collect kinematic data at 100 Hz. Marker clusters were placed bilaterally on the lower back, thigh, shank, and foot. We digitized the joint centers and the posterior aspect of heels to determine the position of heel relative to the marker clusters. Additionally, we digitized the top edge of the obstacle to track its position.

**Figure 1.**
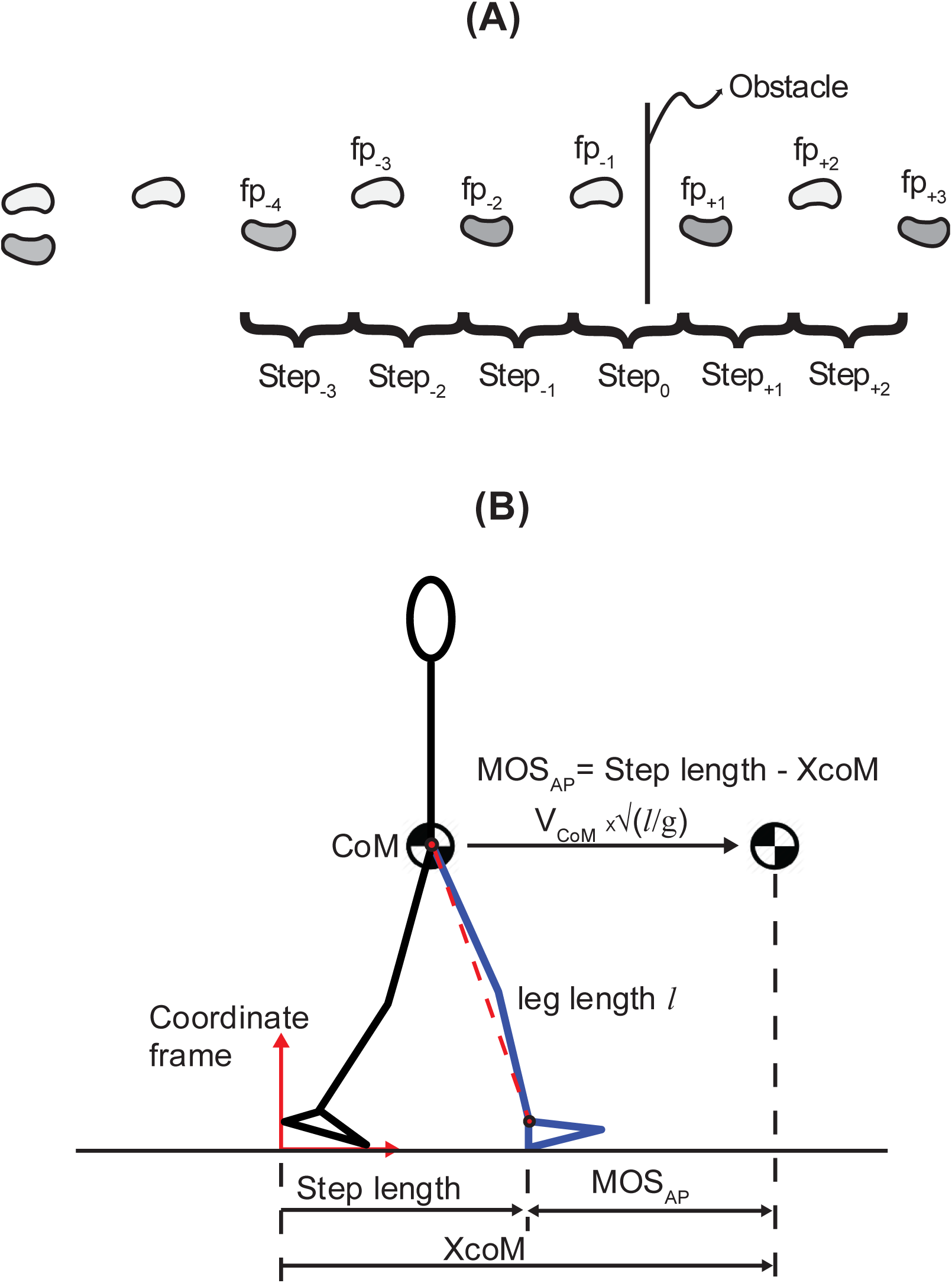
Experimental setup. Panel A shows the walkway layout with foot placements (fp_-4_ to fp_+3_) and steps (step_-3_ to step_+2_) before and after the obstacle. Panel B illustrates XcoM, step length, MOS_AP_ and their relationship.

### 2.3 Analysis

We discarded some trials due to poor kinematic tracking and selected 15 trials with good kinematic data for each participant and walkway to ensure consistent analysis across the participants. All kinematic data were filtered using a zero-lag, 4th order, low-pass Butterworth filter with a 7 Hz cut-off frequency. We identified seven foot placements (fp_-4_ to fp_+3_) using the anteroposterior (AP) position of the heel [19] (Fig. 1A).

Center of mass position (CoM), center of mass velocity (V_CoM_) and MOS_AP_ at heel contact were quantified at the seven foot placements (fp_-4_ to fp_+3_); step length was quantified at six steps (step_-3_ to step_+2_), and the uncontrolled manifold (UCM) analysis was conducted for the six steps.

Step length was determined as the distance between successive heel contacts. The CoM position was estimated as the centroid of the triangle formed by the left and right anterior superior iliac spines and the midpoint between the left and right posterior superior iliac spines [20]. CoM velocity was obtained by differentiating the CoM position. In the sagittal plane, the extrapolated center of mass (XcoM) was computed as:

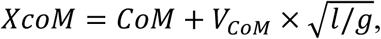

where *g* is the gravitational acceleration, and *l* is the participant’s leg length defined as the sagittal-plane distance between the CoM and the ankle of the limb that touched the ground (Fig. 1B), averaged over the same step across 15 trials [21]. MOS_AP_ was computed at the instant of leading heel contact, in a coordinate frame placed at the location of the rear heel contact (Fig. 1B):

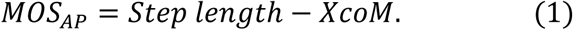

Thus, MOS_AP_ is the distance from the leading heel (anterior boundary of the base of support, BOS) to the XcoM. When MOS_AP_ is negative, XcoM extends beyond the anterior boundary of the BOS (Fig. 1B), and the body possesses sufficient momentum to rotate past vertical and fall forward, assuming that energy loss at heel contact is recovered via an equivalent energy input during push-off. In contrast, a positive MOS_AP_ value indicates that the XcoM is behind the anterior BOS boundary; the body will not rotate beyond the vertical and will fall backward.

MOS_AP_ is a measure of anterior passive stability, appropriate for our purpose of studying stability loss in the forward direction after tripping. Lower, more negative MOS_AP_ (more anterior XcoM) indicates reduced stability, as forward falls become more probable if a person trips. Conversely, larger MOS_AP_ values (more posterior XcoM) indicate enhanced anterior stability [22, 23]. Additionally, MOS_AP_ inversely relates to walking efficiency: lower MOS_AP_ values indicate greater passive forward motion, reducing the active propulsion needed for continued progression, and vice versa. Thus, MOS_AP_ effectively captures the balance between stability and efficiency.

Note that our interpretation of MOS_AP_ is opposite to that implied by Hof [7]. There, the condition MOS_AP_ < 0, when the XcoM projects ahead of the BOS, ensures forward progression. Conversely, when MOS_AP_ > 0, the passive walker’s forward progression ceases. Therefore, a passive walker has stable gait if MOS_AP_ < 0 [7]. We deviate from this result because we want to identify how humans protect themselves against falling forward rather than how they maintain forward progression.

### 2.4 The uncontrolled manifold (UCM) analysis

We used the uncontrolled manifold (UCM) analysis to determine whether step length and XcoM co-vary to stabilize MOS_AP_ at heel contact. The UCM framework quantifies how variability in input variables is structured to maintain stability in a task-relevant output variable. This method has been widely applied in gait research to assess synergy-based control strategies [24, 25].

Briefly, the across-trial variability in the step length and the XcoM for a particular step are partitioned into two subspaces, namely, the UCM and its orthogonal complement (ORT). The UCM is the null space of the Jacobian that relates small changes in step length and XcoM to changes in MOS_AP_. Jacobian (J) is obtained as the partial derivatives of Equation (1): J = [1 −1]. The projection of the deviations in step length and XcoM from their corresponding across-trial means onto the UCM does not affect MOS_AP_, whereas the projection onto the ORT does. Variance in the projections, called V_UCM_ and V_ORT_, respectively, are computed and then combined into the synergy index:

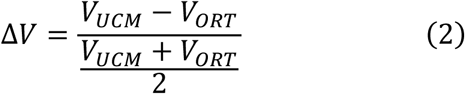

When V_UCM_ > V_ORT_, ΔV > 0, indicating that the step length and XcoM co-vary to stabilize MOS_AP_. Conversely, ΔV < 0 indicates that the inputs covary to change or destabilize MOS_AP_. ΔV = 0 indicates no task-specific co-variation, and that MOS_AP_ is not a controlled variable. The synergy index is bounded within the range of −2 to 2. Therefore, we z-transform ΔV to obtain ΔVz for statistical comparisons [26]:

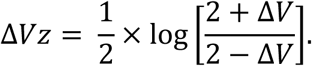

Since ΔV = 0 corresponds to ΔVz = 0, we report ΔVz values in the Results section and use them for statistical inference [24].

### 2.5 Statistics

To assess whether the walkway, foot placement, and age affected MOS_AP_ (H1), we performed a three-way (walkway × step × age) mixed-model ANOVA with walkway and step as repeated measures and age as a between-subjects factor. To identify the source of changes in the MOS_AP_, we applied the same ANOVA separately to the CoM position, CoM velocity at heel contact, and step length.

To assess the presence of synergies (H2), we conducted one-sample t-tests to determine if ΔVz significantly differed from zero for each step on both walkways, for young and older adults.

To assess whether the walkway, foot placement, and age affected the synergy index (H3), we performed a three way (walkway × step × age) mixed-model ANOVA with walkway and step as repeated measures and age as a between-subjects factor. To identify the source of changes in the synergy index, we applied the same ANOVA separately to the variance components (V_UCM_, V_ORT_).

For all ANOVA tests, we fitted a generalized linear model with random effects. Since the UCM analysis yields a single value each for the synergy index, V_UCM_, and V_ORT_, participant was the random effect for these ANOVAs. For all other variables, we used data from all 15 trials, with trial number within each participant as the nested random effect. If no interactions were detected, the significant main effects were followed by pairwise comparisons with Tukey-Kramer adjustments. If interaction effects were present, planned pairwise comparisons with Tukey-Kramer adjustments were conducted to test the hypotheses.

If a three-way interaction was observed, the dependent variable was compared across one independent variable while holding the other two constant. For example, the dependent variable was compared across age groups, with results examined separately for each foot placement and each walkway. If a two-way interaction was observed, pairwise comparisons were planned with the same principle. For example, if the walkway × step interaction was observed, then the dependent variable was compared across walkways at each step (or foot placement), and across-step (or foot placement) for each walkway. All analyses were conducted using the PROC GLIMMIX procedure in SAS 9.4 (Cary, NC, USA) with significance set at 0.05.

## 3. Results

All planned pairwise comparisons are provided in the Appendix. Of the planned pairwise comparisons across foot placements/steps, only neighboring pairs on the obstructed walkway are described below and depicted in Figs. 2 and 3, since they address our hypotheses and keep this section brief.

**Figure 2.**
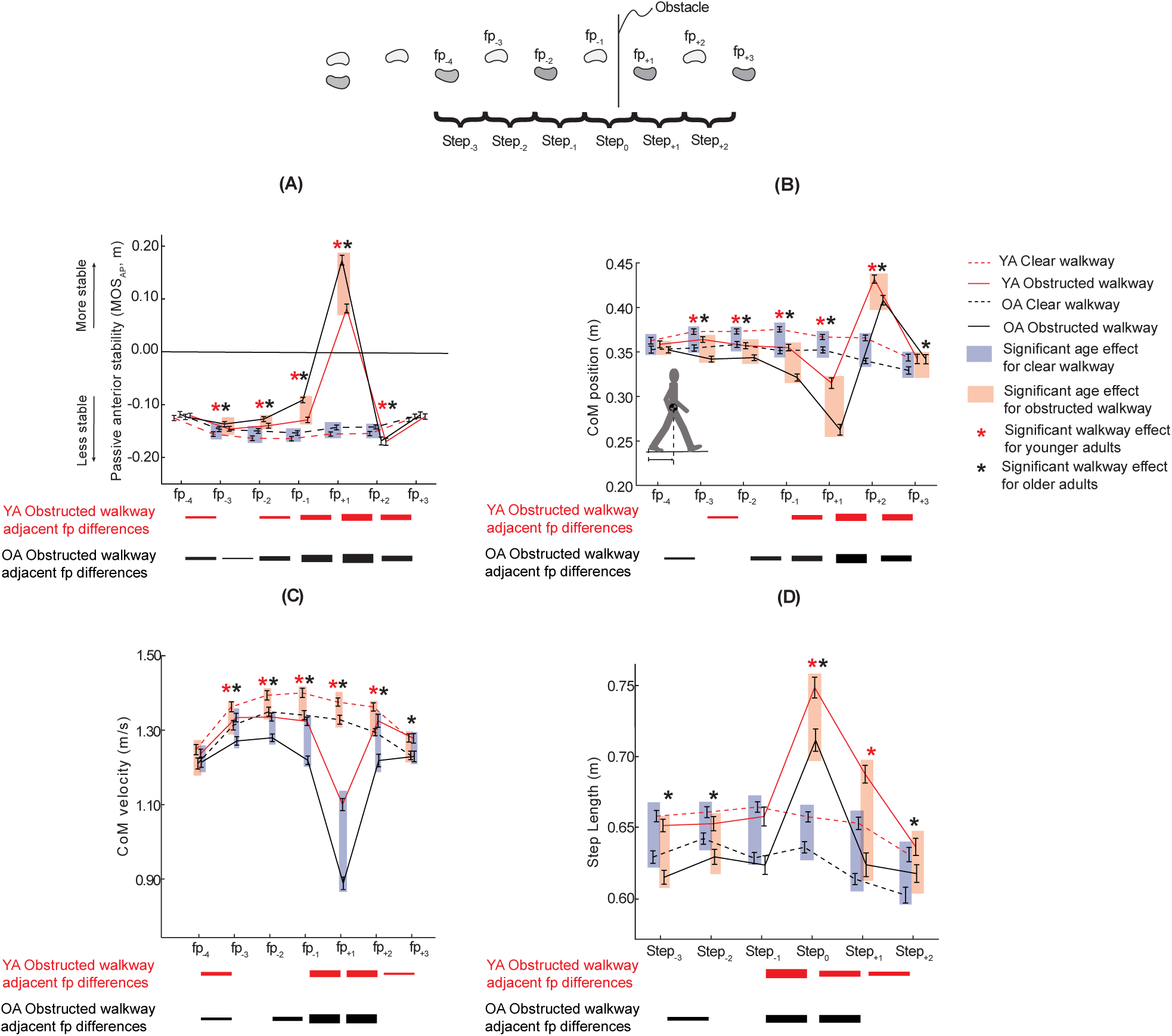
Mean ± standard error for (A) MOS_AP_, (B) CoM position at heel contact, (C) CoM velocity at heel contact, and (D) Step length. Younger adults: red lines; Older adults: black lines; Clear walkway: dashed line; Obstructed walkway: solid line. Shaded box denotes significant age effect, and asterisks indicate significant walkway effect at each foot placement. Solid dashes below each plot indicate significant differences across consecutive foot placements on the obstructed walkway, with line width proportional to estimated mean difference.

**Figure 3.**
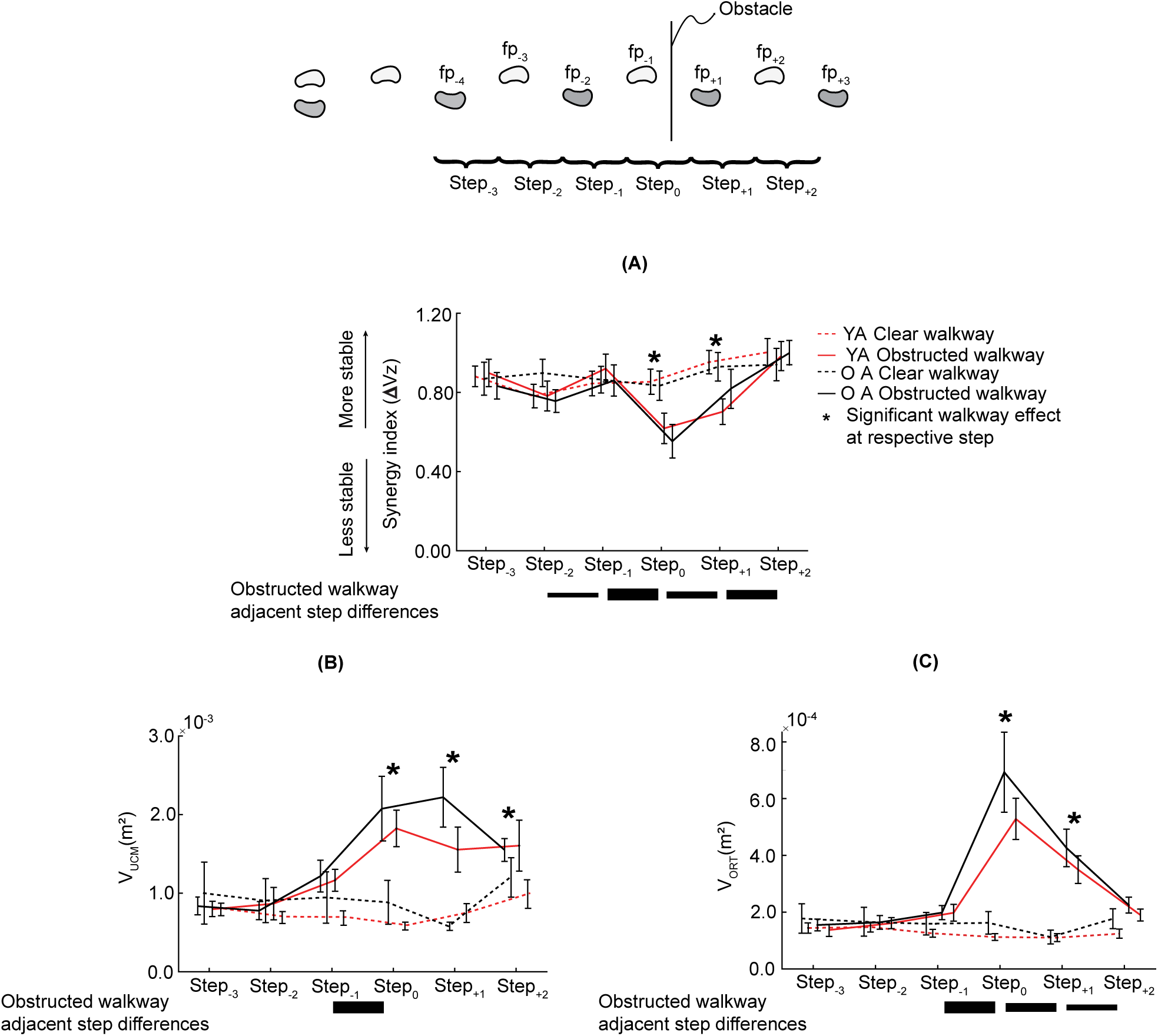
Mean ± standard error for (A) Synergy index, (B) V_UCM_, and (C) V_ORT_. Younger adults: red lines; Older adults: black lines; Clear walkway: dashed line; Obstructed walkway: solid line. Asterisks indicate significant walkway effect at each step. Solid dashes below each plot indicate significant differences across consecutive steps on the obstructed walkway, with line width proportional to estimated mean difference.

### 3.1 Margin of stability (MOS_AP_) and its components

#### MOS_AP_

The across-step pattern of MOS_AP_ was affected by the presence of the obstacle, and the effect was different across the two age groups. This was reflected in a walkway × foot placement × age interaction for MOS_AP_ (F_6,10052_ = 28.34, p < 0.001; η_p_ ^2^ = 0.12; Fig. 2A).

Post hoc comparisons across age revealed that older adults had higher MOS_AP_ compared to younger adults at all foot placements (p < 0.04) with the following exceptions: (1) first and last (fp_-4_ and fp_+3_) in the clear walkway and (2) first (fp_-4_) and the last two (fp_+2_, fp_+3_) in the obstructed walkway (p > 0.09).

Post hoc comparison across walkway revealed similar changes in both age groups. MOS_AP_ was higher in the obstructed walkway from fp_-3_ to fp_+1_ and reversed to a lower value than the clear walkway at fp_+2_ (p < 0.03). Although this pattern was the same in both groups, the estimated across-walkway mean difference was larger for the older adults at fp_-1_, fp_+1_ and fp_+2_, and similar in the two groups for other foot placements.

Post hoc comparisons across consecutive foot placements in the obstructed walkway revealed that MOS_AP_ evolved over multiple steps in both groups. Starting at fp_-3_, MOS_AP_ at each foot placement was different from the preceding one for all foot placements (p < 0.04) except fp_-2_ for young adults (p = 0.13). Although the pattern was the same in both groups, the estimated across-step mean differences were larger in older adults for the two obstacle-crossing steps, namely, transitions from fp_-1_ to fp_+1_ and fp_+1_ to fp_+2_.

#### CoM position relative to rear heel

The across-step pattern of CoM position relative to rear heel was affected by the presence of the obstacle, and the effect was different across the two age groups. This was reflected in walkway × foot placement × age interaction (F_6,10052_ = 13.04, p < 0.001; η_p_^2^ = 0.08; Fig. 2B).

Post hoc comparisons across age revealed that older adults leaned further back (CoM closer to rear heel) compared to younger adults at all foot placements (p < 0.04) except the last (fp_+3_; p = 0.94) in the obstructed walkway.

Post hoc comparisons across walkways revealed that both younger and older adults leaned further back (CoM closer to the rear heel) while approaching and crossing the obstacle with the lead foot (fp_-3_ to fp_+1_) compared to the corresponding steps in the clear walkway (p < 0.01). However, the CoM was farther from the rear heel when the trailing heel contacted the ground (fp_+2_) compared to the corresponding step in the clear walkway (p < 0.01). Although this pattern was the same in both groups, the estimated across-walkway mean difference was larger for the older adults at fp_-1_, fp_+1_ and fp_+2_, and similar in both groups for other foot placements.

Post hoc comparisons across consecutive foot placements in the obstructed walkway revealed that CoM position evolved over multiple steps in both groups. Starting at fp_-3_, older adults changed their CoM position for every foot placement (p < 0.01) except fp_-2_ (p = 0.62). Young adults changed CoM position for every foot placement (p < 0.02) except fp_-3_ and fp_-1_ (p > 0.09). Both groups gradually leaned further back while approaching the obstacle; but the largest changes occurred for the lead and trail crossing foot placements as both groups first rocked back and then forward at fp_+1_ and fp_+2_. Although the pattern was the same in both groups, the estimated across-step mean differences were larger in older adults for the two obstacle-crossing steps.

#### CoM velocity at heel contact

The across-step pattern of CoM velocity was affected by the presence of the obstacle, and the effect was different across the two age groups. This was reflected in walkway × foot placement × age interaction for CoM velocity at heel contact (F_6,10052_ = 9.61, p < 0.001; η_p_ ^2^ = 0.07; Fig. 2C).

Post hoc comparisons across age revealed that older adults had lower CoM velocity compared to younger adults at all foot placements (p < 0.01) except the first (fp_-4_) on the obstructed walkway (p = 0.09).

Post hoc comparisons across walkways revealed that both young and older adults had lower CoM velocity at all foot placements (p < 0.03) except the first and last (fp_-4_, fp_+3_; p > 0.41) in the obstructed compared to the clear walkway. Although this pattern was the same in both groups, the estimated across-walkway mean difference was larger for the older adults at all foot placements that demonstrated an age effect.

Post hoc comparisons across consecutive foot placements on the obstructed walkway revealed that CoM velocity evolved over multiple approach steps. Both groups increased velocity from fp_-4_ to fp_-3_ (p < 0.01). However, the older adults initiated velocity reduction at fp_-1_, whereas young adults first reduced velocity at fp_+1_ (p < 0.01). The changes for the lead and trail crossing foot placements were large, as both groups first slowed down and then sped up at fp_+1_ and fp_+2_ (p < 0.01). For the two obstacle-crossing steps, although the pattern was the same in both groups, the estimated across-step mean differences were larger in older adults.

#### Step length

The across-step pattern of step length was affected by the presence of the obstacle, and the effect was different across the two age groups. This was reflected in walkway × foot placement × age interaction for step length (F_6,8616_ = 2.65, p = 0.02; η_p_ ^2^ = 0.03; Fig. 2D).

Post hoc comparisons across age revealed that older adults had shorter step length compared to younger adults at all steps for both walkways (p < 0.01).

Post hoc comparisons across walkways revealed that older adults took a longer step while crossing the obstacle with the lead foot (step_0_) but their steps during approach (step_-3_, step_-2_) were shorter compared to the corresponding steps in the clear walkway (p < 0.02). Young adults also took longer steps while crossing (step_0_, step_+1_; p < 0.01), but unlike the older adults, their step length during approach was not different on the two walkways (p > 0.11). The estimated across-walkway mean difference was smaller for the older adults for the obstacle-crossing steps.

Post hoc comparisons across consecutive steps on the obstructed walkway revealed slight increase in step length while approaching the obstacle for older adults. The major changes were for the two crossing steps in both groups (step_0_, step_+1_; p < 0.01). The magnitude of step length changes was similar, but the pattern was different in the two groups. Lead step length (step_0_) was longer than preceding steps in both age groups. For older adults, trail step length (step_+1_) was not different from approach steps. However, for younger adults, it was longer than approach steps, but shorter than lead step length.

### 3.2 UCM variables

#### Synergy Index

Participants covaried their step length and XcoM to stabilize MOS_AP_ for each step of both walkways. This was reflected in positive synergy index (ΔVz) for all steps regardless of age (young adults: t_24_ ≥ 8.96, p < 0.01, Cohen’s d ≥ 2.98; older adults: t_22_ ≥ 7.00; p < 0.01; Cohen’s d ≥ 2.90; Fig. 3A).

The across-step pattern of the synergy index was affected by the obstacle, particularly at the crossing step and the following step. This was reflected in a walkway × step interaction for ΔVz (F_5,506_ = 3.75, p = 0.002; η_p_ ^2^ = 0.03; Fig. 3A).

Post hoc comparisons across walkways revealed that ΔVz was lower on the obstructed compared to the clear walkway for both the lead (step_0_) and trail crossing steps (step_+1_).

Post hoc comparisons across consecutive steps on the obstructed walkway revealed that ΔVz increased at one approach step (step_-1_), declined for the lead crossing step (step_0_) and recovered over the next two steps.

#### V_UCM_

The across-step pattern in V_UCM_ was affected by the presence of the obstacle. This was reflected in a walkway × step interaction for V_UCM_ (F_5,506_ = 8.77, p < 0.0001; η_p_ ^2^ = 0.07; Fig. 3B).

Post hoc comparisons across walkways revealed that V_UCM_ was higher on the obstructed compared to the clear walkway at all steps except the first two (step_-3_ and step_-2_).

Post hoc comparisons across consecutive steps on the obstructed walkway revealed that V_UCM_ increased for the lead crossing step (step_0_).

#### V_ORT_

The across-step pattern in V_ORT_ was affected by the presence of the obstacle. This was reflected in a walkway × step interaction for V_ORT_ (F_5, 506_ = 25.31, p < 0.0001; η_p_ ^2^ = 0.10; Fig. 3C).

Post hoc comparisons across walkways revealed that V_ORT_ was higher on the obstructed compared to the clear walkway at the lead (step_0_) and trail (step_+1_) crossing steps.

Post hoc comparisons across consecutive steps on the obstructed walkway revealed that V_ORT_ increased for the lead crossing step (step_0_) and then recovered over the next two steps.

## 4. DISCUSSION

We report two main findings: (1) humans actively recruit passive mechanics as an insurance policy to protect against trips turning into falls, as reflected in increasing MOS_AP_ and positive synergy indices during obstacle crossing; and (2) aging affects MOS_AP_ but does not alter the synergy index. Recall the key distinction in this work: mean MOS_AP_ captures the passive dynamic stability of gait arising from the physics of body motion, whereas the synergy index quantifies the active neural coordination of XcoM position and foot placement to stabilize MOS_AP_ at specific values.

As predicted in H1, both groups increased MOS_AP_ from step-to-step on the obstructed walkway, with older adults demonstrating greater stability in both walkways and greater increase in MOS_AP_ during obstacle crossing. As predicted in H2, both age groups corrected step length in response to XcoM deviations, stabilizing MOS_AP_ at a specific value at each heel contact. The synergy index was consistently positive across all participants and conditions, demonstrating active neural coordination of XcoM position and foot placement. Finally, hypothesis H3 was partially supported. As expected, the synergy index was lower for the crossing steps than the corresponding steps in unobstructed gait, but it did not weaken while approaching the obstacle. Age did not influence this pattern.

### 4.1 Active MOS_AP_ Control

Control of obstructed gait in the AP direction is active, relying on supraspinal processes that modulate motor action based on integrated sensory information, and manifests as control of MOS_AP_. This idea has some precedent. Hof demonstrated that a consistent MOS_AP_ yields stable gait in a mathematical model and suggested that humans might employ this control strategy [7]. The idea has since been implicitly invoked to explain deviations in step length and/or speed in response to perturbations. The deviations that arise when adults are forced to walk with shorter steps or at higher speeds [23] and during corrective movements after transient mechanical perturbations [27] yield systematic changes in MOS_AP_. These changes are interpreted as adaptations to increase passive dynamic stability for reducing fall risk, with changes in foot placements as the underlying mechanism.

We contribute here by providing direct evidence of active MOS_AP_ control. Strong evidence comes from the systematic increase in MOS_AP_ during approach and crossing, contrasting with the relatively uniform pattern observed during unobstructed walking. MOS_AP_ reaches its peak during the lead crossing step, when the risk of falling from a trip is greatest [28]. A higher MOS_AP_ indicates a more posterior XcoM and reduced forward momentum at heel contact, which suggests lower forward momentum at the preceding swing phase when the foot crosses the obstacle. This reduced momentum lowers the risk that a trip will result in a fall.

Further support for this interpretation is that while both groups increased MOS_AP_ for the obstacle-crossing steps (fp_-1_, fp_+1_), older adults amplified the increase relative to younger adults. Age related deficits in cognitive and sensory-motor capabilities [3] increase response times and limit amplitudes of corrective neuromuscular responses after a trip [4]. Furthermore, falls carry more severe consequences for older adults [29]. Older adults adapt by increasing their anterior passive stability, relying more on passive dynamics for protection, thereby pointing to active control of MOS_AP_.

We next consider how this control is achieved. MOS_AP_ rises when the XcoM moves backward either because the CoM moves backwards (backward lean) and/or slows, or when step length increases, moving the BOS boundary forward. During unobstructed walking, speed, step length, and lower-limb joint powers are empirically coupled: faster walking is accompanied by longer steps [30] and greater lower-limb joint power without redistribution of relative joint contributions [31]. During approach (i.e., fp_-3_ to fp_-1_, since CoM velocity was rising from fp_-4_ to fp-_3_ from gait initiation; Fig. 2C), mean MOS_AP_ rose mainly through CoM motion: backward lean and lower speed, while mean step length stayed flat (Fig. 2D). At step_0_, the step with the greatest fall risk, the CoM slowed further and step length increased. This pattern violates the empirical coupling evident during unobstructed walking and therefore cannot be interpreted as its speed-scaled version. The pattern is consistent with task-specific regulation of foot placement and reorganization of limb kinetics, including altered push-off and swing-phase kinetics [32]. This interpretation is supported by the UCM analysis, which independently identified step length as the variable compensating for trial-to-trial deviations in XcoM, holding MOS_AP_ near its mean throughout approach and crossing (V_UCM_ > V_ORT_; positive synergy index; Fig. 3).

### 4.2 Passive Stabilization vs. Supraspinal Control

The increase in mean MOS_AP_ and the positive synergy index arise from three sources: passive mechanics, spinal feedback, and supraspinal control. During unobstructed walking, the sagittal-plane gait cycle is passively stable: step length scales with walking speed, and deviations in speed produce proportional step-length changes through the passive pendular motion of the swinging leg, restoring the cycle to its mean. The process is assisted by spinal feedback, but operates without supraspinal influence [12, 30, 33]. Most step-length variability on clear walkways is attributable to slow, self-selected fluctuations in speed rather than corrective action [34]. This passive process alone generates a positive correlation between CoM speed and step length, and with it, a degree of covariation between XcoM and step length that stabilizes MOS_AP_ about whatever average it happens to take.

On obstructed walkways, however, supraspinal control intervenes. It modulates behavior while approaching and navigating hazards. Humans direct their gaze to footholds two steps ahead to update body state before each step [35], change walking speed two steps before an even-to-uneven surface transition [36], and sample an obstacle’s location and dimensions during approach to plan foot placement and limb elevation [37]. Our work identifies preparatory adjustments up to two steps before the obstacle, evident in MOS_AP_ and its components (Fig. 2). Processing visual information and issuing timed motor commands are supraspinal functions; EEG recordings confirm that prefrontal activity rises during preparation for the crossing step and adjusts spinal reflex gain in the stance limb beforehand [38]. Supraspinal control influences the mean speed, body posture, and step length, and through them, sets the mean MOS_AP_ that passive dynamics and spinal feedback then stabilize about.

The synergy index, therefore, captures the combined output of three processes: passive covariation between CoM speed and step length, spinal feedback, and supraspinal tuning that sets the gain and target for that feedback. It cannot partition their separate contributions, a property of synergies generally [39]. The cleaner signature of active control is therefore the systematic, anticipatory change in mean MOS_AP_, a pattern with no passive explanation, whereas the synergy analysis provides supporting evidence and helps identify the mechanisms involved.

### 4.3 Energetic Cost of Stability

The increased protection against falls provided by a high MOS_AP_ comes at the cost of efficiency: an energetic premium. This is clear for the lead crossing step (step_0_) that pairs the lowest CoM velocity with the longest step length across all steps, a combination that violates well-known speed-step-length optimum. Humans vary step length systematically with walking speed to minimize the combined cost of step-to-step collision work and leg swing [30]. Since the step length for step_0_ is longer than its speed warrants, the rate of negative work required to redirect the CoM velocity during the step-to-step transition, which scales with the fourth power of step length [40], is disproportionately elevated, and the trailing limb must produce disproportionally greater push-off work to compensate. Step_0_ thus incurs a disproportionately high collision-and-push-off penalty.

Apart from the crossing step, energetic expense is likely higher while approaching the obstacle, but the evidence is less direct. During approach, MOS_AP_ increased via a posterior shift in CoM position. When the CoM sits further behind the stance ankle, the trailing leg must push off harder to redirect the CoM over and past the stance ankle and sustain forward progression. Backward trunk lean, which shifts the CoM posteriorly, has been shown to increase push-off work and metabolic cost [41], suggesting that a similar energetic penalty likely occurred during the approach phase.

### 4.4 Selective Preservation of Synergies with Aging

We have reported three important age-related patterns. First, the obstacle has an amplified effect on MOS_AP_ in older adults, which we interpreted as intelligent, adaptive deployment of the body’s mechanics to minimize fall risk. Second, age did not affect the synergy index. Third, older adults initiated changes earlier on the obstructed walkway.

Aging should weaken synergies because it impacts two things that synergies require: neural coupling among the body’s abundant degrees of freedom (DOF; muscle fibers, joints, segments, etc.), and reliable sensory information about body state and task variables [39]. During development, humans learn to build task-specific couplings among DOF and recruit them flexibly to stabilize important motor outcomes. Aging degrades the neural and sensory processes that support these couplings [3]. The system should be less able to exploit the abundant DOF. Control should become more element-based, with fewer compensatory covariations, lower V_UCM_, higher V_ORT_, and weaker synergies [25].

This principled prediction is not supported by the data; healthy aging does not produce a global weakening of locomotor synergies. Some studies report weaker synergies [17, 26, 42], others report preserved synergies [16, 43, 44], and some even report stronger synergies in older adults [45, 46]. Importantly, these studies quantify synergies stabilizing different kinematic and kinetic locomotor variables. This suggests one explanation: healthy older adults reorganize control to preserve synergies for variables most directly tied to balance, while tolerating weaker stabilization of less critical gait variables. Across locomotor tasks, mediolateral swing-foot trajectory, mediolateral CoM position, net ground reaction forces and moments, leading-limb toe clearance, and, based on the present data, MOS_AP_, belong to the balance-critical class, since synergies stabilizing these variables do not weaken with age. In contrast, trailing-limb toe height and step length during obstacle crossing may be less critical to balance, since synergies stabilizing these variables weaken with age. The increased variability in these quantities does not immediately compromise dynamic equilibrium in older adults; it is easier to recover from a trail trip [28], and step length is subservient to a more important gait variable – MOS_AP_.

As noted above, older adults initiated changes earlier on the obstructed walkway. MOS_AP_ rises and backward lean and CoM speed change one step earlier compared to young adults (Fig. 2). This could be an adaptation to the general slowing with age, further highlighting intelligent deployment of available resources.

### 4.5 Limitations

The main limitations of this study arise from the stability measure. MOS is a physicist’s spherical cow, a deliberately simple model that captures some essential features of locomotion at the cost of biological complexity. The underlying model is the passive inverted pendulum with the body modeled as a point mass atop a rigid leg, which ignores neuromuscular control, trunk angular momentum, compliant joints, etc. Three limitations follow from this simplification.

First, MOS_AP_ quantifies only passive dynamic stability. This is a problem because gait stability emerges from passive dynamics and active neuromuscular control, and MOS_AP_ ignores the latter. As explained earlier, positive MOS_AP_ at heel strike, where XcoM is behind the anterior BOS boundary, indicates that a passive walker does not have sufficient energy to rotate past vertical and it will stall. However, a human could compensate by increasing push-off work from the rear leg. Indeed, positive MOS_AP_ has been observed in stable human walking [9], and both positive and negative values appeared for both age groups in our data. MOS_AP_ therefore does not provide a threshold that classifies gait as stable or unstable, a limitation acknowledged for several biomechanical stability measures [47]. Rather, it provides a graded, continuous index that yields meaningful information only with appropriate context.

Second, MOS_AP_ ignores the head, arms, and trunk. The foot placement estimator extends the inverted pendulum logic to a model with a trunk possessing mass and moment of inertia and provides a conceptually analogous but more complete stability measure [48]. We expect our core findings to hold under this more complex model, but this remains to be tested.

Third, we did not measure metabolic energy consumption directly. The stability-efficiency tradeoff was inferred from physics-based arguments and the data. Direct measurement of metabolic cost during obstacle negotiation is a natural and important next step.

## 5. Conclusion

Our data indicate that human adults actively recruit body mechanics, adjusting posture and momentum so that the body’s passive response to a potential trip acts more in favor of recovery than a forward fall. Persistent XcoM–step length covariation confirms that this mechanical reorganization is active and continuous across approach and crossing steps. Older adults amplify this strategy to compensate for diminished neuromuscular corrective capabilities, a rather covert adjustment, unlike the overt use of external aids. This anticipatory mechanical reorganization is not confined to skilled athletes; healthy walking adults across the lifespan show it during routine movements. This is embodied natural intelligence.

## Funding

AK was funded by Bilsland Dissertation Fellowship, Purdue University

## Competing Interests

The authors declare that they have no competing interests.

## Data availability

Data and code are published by the Purdue University Research Repository and are publicly available for download [49].

## Authors’ contributions

A.K.: data curation, formal analysis, investigation, methodology, project administration, resources, software, validation, visualization, writing-original draft, writing—review and editing; C.C.: investigation, writing-review and editing; S.R.: Investigation, project administration, resources, writing-review and editing; S.A.: Conceptualization, investigation, methodology, project administration, resources, writing-original draft, writing-review and editing.

All authors gave final approval for publication and agreed to be held accountable for the work performed therein.

## AI statement

AI was used solely to improve readability and language of this manuscript.

## Notes

https://purr.purdue.edu/publications/4150/usage?v=2

### Competing Interest Statement

The authors have declared no competing interest.

