## Appendix for "Older adults amplify passive gait stability during obstacle crossing without weakening the stabilizing synergy"

**Appendix A**

Pre-planned pairwise comparisons of simple effects following the significant age × walkway × foot placement interaction for $MOS_{AP}$. Each comparison tested one independent variable while holding the other two constant. Thus, age-group comparisons were evaluated separately for each walkway and foot placement; walkway comparisons were evaluated separately for each age group and foot placement; and foot placement comparisons were evaluated separately for each age group and walkway. Least-squares mean differences are reported with estimates, degrees of freedom, t values, and p values. $MOS_{AP}$ = margin of stability; OA = older adults; YA = younger adults; df = degrees of freedom.

**Table 1.1. Type III tests of fixed effects for $MOS_{AP}$.**

| Effect | Num DF | Den DF | F value | Pr > F |
|---|---|---|---|---|
| Age | 1 | 10052 | 197.17 | <.0001 |
| Walkway | 1 | 10052 | 1834.25 | <.0001 |
| Age × Walkway | 1 | 10052 | 26.32 | <.0001 |
| fp | 6 | 10052 | 1095.04 | <.0001 |
| Age × fp | 6 | 10052 | 33.00 | <.0001 |
| Walkway × fp | 6 | 10052 | 1160.94 | <.0001 |
| Age × Walkway × fp | 6 | 10052 | 28.34 | <.0001 |

**Table 1.2. Pairwise comparisons between clear and obstructed walkway for least-squares mean $MOS_{AP}$, evaluated separately for each age group and foot placement.**

| Simple Effect Level | | Walkway | Walkway | Estimate | df | t value | Pr > \|t\| |
|---|---|---|---|---|---|---|---|
| OA | $fp_{-4}$ | Clear walkway | Obstructed walkway | 0.005 | 10052 | 1.09 | 0.27 |
| OA | $fp_{-3}$ | Clear walkway | Obstructed walkway | -0.010 | 10052 | -2.30 | 0.02* |
| OA | $fp_{-2}$ | Clear walkway | Obstructed walkway | -0.022 | 10052 | -5.16 | <.0001* |
| OA | $fp_{-1}$ | Clear walkway | Obstructed walkway | -0.062 | 10052 | -14.31 | <.0001* |
| OA | $fp_{+1}$ | Clear walkway | Obstructed walkway | -0.316 | 10052 | -72.61 | <.0001* |
| OA | $fp_{+2}$ | Clear walkway | Obstructed walkway | 0.0265 | 10052 | 6.08 | <.0001* |
| OA | $fp_{+3}$ | Clear walkway | Obstructed walkway | -0.003 | 10052 | -0.70 | 0.48 |
| YA | $fp_{-4}$ | Clear walkway | Obstructed walkway | -0.005 | 10052 | -1.11 | 0.27 |
| YA | $fp_{-3}$ | Clear walkway | Obstructed walkway | -0.009 | 10052 | -2.32 | 0.02* |
| YA | $fp_{-2}$ | Clear walkway | Obstructed walkway | -0.023 | 10052 | -5.68 | <.0001* |
| YA | $fp_{-1}$ | Clear walkway | Obstructed walkway | -0.035 | 10052 | -8.49 | <.0001* |
| YA | $fp_{+1}$ | Clear walkway | Obstructed walkway | -0.237 | 10052 | -56.84 | <.0001* |
| YA | $fp_{+2}$ | Clear walkway | Obstructed walkway | 0.015 | 10052 | 3.70 | 0.0002* |
| YA | $fp_{+3}$ | Clear walkway | Obstructed walkway | -0.005 | 10052 | -1.31 | 0.19 |

**Table 1.3. Pairwise comparisons between younger and older adults for least-squares mean $MOS_{AP}$, evaluated separately for each walkway and foot placement.**

| Simple Effect Level | Age | Age | Estimate | DF | t Value | Pr > \|t\| |
|---|---|---|---|---|---|---|
| Clear walkway fp-4 | OA | YA | 0.007 | 10052 | 1.66 | 0.09 |
| Clear walkway fp-3 | OA | YA | 0.009 | 10052 | 2.14 | 0.03* |
| Clear walkway fp-2 | OA | YA | 0.014 | 10052 | 3.21 | <0.01* |
| Clear walkway fp-1 | OA | YA | 0.010 | 10052 | 2.51 | 0.01* |
| Clear walkway fp+1 | OA | YA | 0.012 | 10052 | 2.91 | <0.01* |
| Clear walkway fp+2 | OA | YA | 0.012 | 10052 | 2.94 | <0.01* |
| Clear walkway fp+3 | OA | YA | 0.005 | 10052 | 1.31 | 0.19 |
| Obstructed walkway fp-4 | OA | YA | -0.002 | 10052 | -0.54 | 0.59 |
| Obstructed walkway fp-3 | OA | YA | 0.009 | 10052 | 2.22 | 0.03* |
| Obstructed walkway fp-2 | OA | YA | 0.012 | 10052 | 2.91 | <0.01* |
| Obstructed walkway fp-1 | OA | YA | 0.037 | 10052 | 8.80 | <.0001* |
| Obstructed walkway fp+1 | OA | YA | 0.091 | 10052 | 21.38 | <.0001* |
| Obstructed walkway fp+2 | OA | YA | 0.001 | 10052 | 0.36 | 0.72 |
| Obstructed walkway fp+3 | OA | YA | 0.003 | 10052 | 0.74 | 0.46 |

**Table 1.4. Pairwise comparisons between foot placements for least-squares mean $MOS_{AP}$, evaluated separately for each age group and walkway.**

| Simple Effect Level | fp | fp | Estimate | DF | t Value | Pr > \|t\| |
|---|---|---|---|---|---|---|
| Age × Walkway OA Clear walkway | $fp_{+1}$ | $fp_{+2}$ | -0.0009 | 10052 | -0.22 | 0.82 |
| Age × Walkway OA Clear walkway | $fp_{+1}$ | $fp_{+3}$ | -0.0207 | 10052 | -4.76 | <.0001* |
| Age × Walkway OA Clear walkway | $fp_{+1}$ | $fp_{-1}$ | 0.0105 | 10052 | 2.42 | 0.01* |
| Age × Walkway OA Clear walkway | $fp_{+1}$ | $fp_{-2}$ | 0.0068 | 10052 | 1.54 | 0.12 |
| Age × Walkway OA Clear walkway | $fp_{+1}$ | $fp_{-3}$ | 0.0034 | 10052 | 0.78 | 0.43 |
| Age × Walkway OA Clear walkway | $fp_{+1}$ | $fp_{-4}$ | -0.0249 | 10052 | -5.72 | <.0001* |
| Age × Walkway OA Clear walkway | $fp_{+2}$ | $fp_{+3}$ | -0.0197 | 10052 | -4.54 | <.0001* |
| Age × Walkway OA Clear walkway | $fp_{+2}$ | $fp_{-1}$ | 0.0114 | 10052 | 2.64 | <0.01* |
| Age × Walkway OA Clear walkway | $fp_{+2}$ | $fp_{-2}$ | 0.00785 | 10052 | 1.78 | 0.07 |
| Age × Walkway OA Clear walkway | $fp_{+2}$ | $fp_{-3}$ | 0.00437 | 10052 | 1.00 | 0.31 |
| Age × Walkway OA Clear walkway | $fp_{+2}$ | $fp_{-4}$ | -0.0239 | 10052 | -5.49 | <.0001* |
| Age × Walkway OA Clear walkway | $fp_{+3}$ | $fp_{-1}$ | 0.0312 | 10052 | 7.18 | <.0001* |
| Age × Walkway OA Clear walkway | $fp_{+3}$ | $fp_{-2}$ | 0.0274 | 10052 | 6.30 | <.0001* |
| Age × Walkway OA Clear walkway | $fp_{+3}$ | $fp_{-3}$ | 0.0241 | 10052 | 5.55 | <.0001* |
| Age × Walkway OA Clear walkway | $fp_{+3}$ | $fp_{-4}$ | -0.0041 | 10052 | -0.95 | 0.34 |
| Age × Walkway OA Clear walkway | $fp_{-1}$ | $fp_{-2}$ | -0.0038 | 10052 | -0.88 | 0.37 |
| Age × Walkway OA Clear walkway | $fp_{-1}$ | $fp_{-3}$ | -0.0071 | 10052 | -1.63 | 0.10 |
| Age × Walkway OA Clear walkway | $fp_{-1}$ | $fp_{-4}$ | -0.0354 | 10052 | -8.13 | <.0001* |
| Age × Walkway OA Clear walkway | $fp_{-2}$ | $fp_{-3}$ | -0.0032 | 10052 | -0.75 | 0.45 |
| Age × Walkway OA Clear walkway | $fp_{-2}$ | $fp_{-4}$ | -0.0315 | 10052 | -7.25 | <.0001* |

| | | | | | | | |
|---|---|---|---|---|---|---|---|
| Age × Walkway | OA Clear walkway | $fp_{-3}$ | $fp_{-4}$ | -0.0283 | 10052 | -6.50 | <.0001* |
| Age × Walkway | OA Obstructed walkway | $fp_{+1}$ | $fp_{+2}$ | 0.3418 | 10052 | 78.47 | <.0001* |
| Age × Walkway | OA Obstructed walkway | $fp_{+1}$ | $fp_{+3}$ | 0.2925 | 10052 | 67.15 | <.0001* |
| Age × Walkway | OA Obstructed walkway | $fp_{+1}$ | $fp_{-1}$ | 0.2845 | 10052 | 60.72 | <.0001* |
| Age × Walkway | OA Obstructed walkway | $fp_{+1}$ | $fp_{-2}$ | 0.3008 | 10052 | 68.99 | <.0001* |
| Age × Walkway | OA Obstructed walkway | $fp_{+1}$ | $fp_{-3}$ | 0.3097 | 10052 | 71.10 | <.0001* |
| Age × Walkway | OA Obstructed walkway | $fp_{+1}$ | $fp_{-4}$ | 0.2962 | 10052 | 67.99 | <.0001* |
| Age × Walkway | OA Obstructed walkway | $fp_{+2}$ | $fp_{+3}$ | -0.0493 | 10052 | -11.32 | <.0001* |
| Age × Walkway | OA Obstructed walkway | $fp_{+2}$ | $fp_{-1}$ | -0.0773 | 10052 | -17.75 | <.0001* |
| Age × Walkway | OA Obstructed walkway | $fp_{+2}$ | $fp_{-2}$ | -0.0412 | 10052 | -9.48 | <.0001* |
| Age × Walkway | OA Obstructed walkway | $fp_{+2}$ | $fp_{-3}$ | -0.0321 | 10052 | -7.37 | <.0001* |
| Age × Walkway | OA Obstructed walkway | $fp_{+2}$ | $fp_{-4}$ | -0.0456 | 10052 | -10.48 | <.0001* |
| Age × Walkway | OA Obstructed walkway | $fp_{+3}$ | $fp_{-1}$ | -0.0280 | 10052 | -6.43 | <.0001* |
| Age × Walkway | OA Obstructed walkway | $fp_{+3}$ | $fp_{-2}$ | 0.0080 | 10052 | 1.84 | 0.06 |
| Age × Walkway | OA Obstructed walkway | $fp_{+3}$ | $fp_{-3}$ | 0.0171 | 10052 | 3.95 | <.0001* |
| Age × Walkway | OA Obstructed walkway | $fp_{+3}$ | $fp_{-4}$ | 0.00365 | 10052 | 0.84 | 0.40 |
| Age × Walkway | OA Obstructed walkway | $fp_{-1}$ | $fp_{-2}$ | 0.0360 | 10052 | 8.27 | <.0001* |
| Age × Walkway | OA Obstructed walkway | $fp_{-1}$ | $fp_{-3}$ | 0.0452 | 10052 | 10.38 | <.0001* |
| Age × Walkway | OA Obstructed walkway | $fp_{-1}$ | $fp_{-4}$ | 0.0316 | 10052 | 7.27 | <.0001* |
| Age × Walkway | OA Obstructed walkway | $fp_{-2}$ | $fp_{-3}$ | 0.0091 | 10052 | 2.10 | 0.03* |
| Age × Walkway | OA Obstructed walkway | $fp_{-2}$ | $fp_{-4}$ | -0.0043 | 10052 | -1.00 | 0.31 |
| Age × Walkway | OA Obstructed walkway | $fp_{-3}$ | $fp_{-4}$ | -0.0135 | 10052 | -3.11 | 0.002* |
| Age × Walkway | YA Clear walkway | $fp_{+1}$ | $fp_{+2}$ | -0.0008 | 10052 | -0.20 | 0.83 |
| Age × Walkway | YA Clear walkway | $fp_{+1}$ | $fp_{+3}$ | -0.0275 | 10052 | -6.60 | <.0001* |
| Age × Walkway | YA Clear walkway | $fp_{+1}$ | $fp_{-1}$ | 0.00882 | 10052 | 2.11 | 0.03* |
| Age × Walkway | YA Clear walkway | $fp_{+1}$ | $fp_{-2}$ | 0.0079 | 10052 | 1.91 | 0.06 |
| Age × Walkway | YA Clear walkway | $fp_{+1}$ | $fp_{-3}$ | 0.0001 | 10052 | 0.03 | 0.97 |
| Age × Walkway | YA Clear walkway | $fp_{+1}$ | $fp_{-4}$ | -0.0302 | 10052 | -7.23 | <.0001* |
| Age × Walkway | YA Clear walkway | $fp_{+2}$ | $fp_{+3}$ | -0.0287 | 10052 | -6.39 | <.0001* |
| Age × Walkway | YA Clear walkway | $fp_{+2}$ | $fp_{-1}$ | 0.0098 | 10052 | 2.31 | 0.02* |
| Age × Walkway | YA Clear walkway | $fp_{+2}$ | $fp_{-2}$ | 0.0088 | 10052 | 2.11 | 0.03* |
| Age × Walkway | YA Clear walkway | $fp_{+2}$ | $fp_{-3}$ | 0.00098 | 10052 | 0.23 | 0.81 |
| Age × Walkway | YA Clear walkway | $fp_{+2}$ | $fp_{-4}$ | -0.0293 | 10052 | -7.03 | <.0001* |
| Age × Walkway | YA Clear walkway | $fp_{+3}$ | $fp_{-1}$ | 0.0363 | 10052 | 8.71 | <.0001* |
| Age × Walkway | YA Clear walkway | $fp_{+3}$ | $fp_{-2}$ | 0.0355 | 10052 | 8.51 | <.0001* |
| Age × Walkway | YA Clear walkway | $fp_{+3}$ | $fp_{-3}$ | 0.0277 | 10052 | 6.63 | <.0001* |
| Age × Walkway | YA Clear walkway | $fp_{+3}$ | $fp_{-4}$ | -0.0026 | 10052 | -0.63 | 0.53 |
| Age × Walkway | YA Clear walkway | $fp_{-1}$ | $fp_{-2}$ | -0.0008 | 10052 | -0.20 | 0.84 |
| Age × Walkway | YA Clear walkway | $fp_{-1}$ | $fp_{-3}$ | -0.0086 | 10052 | -2.08 | 0.04* |
| Age × Walkway | YA Clear walkway | $fp_{-1}$ | $fp_{-4}$ | -0.0390 | 10052 | -9.34 | <.0001* |
| Age × Walkway | YA Clear walkway | $fp_{-2}$ | $fp_{-3}$ | -0.0078 | 10052 | -1.88 | 0.0603 |
| Age × Walkway | YA Clear walkway | $fp_{-2}$ | $fp_{-4}$ | -0.0381 | 10052 | -9.14 | <.0001* |
| Age × Walkway | YA Clear walkway | $fp_{-3}$ | $fp_{-4}$ | -0.0303 | 10052 | -7.26 | <.0001* |
| Age × Walkway | YA Obstructed walkway | $fp_{+1}$ | $fp_{+2}$ | 0.2521 | 10052 | 60.34 | <.0001* |
| Age × Walkway | YA Obstructed walkway | $fp_{+1}$ | $fp_{+3}$ | 0.2044 | 10052 | 48.93 | <.0001* |

| | | | | | | |
|---|---|---|---|---|---|---|
| Age × Walkway YA Obstructed walkway | $fp_{+1}$ | $fp_{-1}$ | 0.2108 | 10052 | 50.45 | <.0001* |
| Age × Walkway YA Obstructed walkway | $fp_{+1}$ | $fp_{-2}$ | 0.2217 | 10052 | 53.07 | <.0001* |
| Age × Walkway YA Obstructed walkway | $fp_{+1}$ | $fp_{-3}$ | 0.2279 | 10052 | 54.55 | <.0001* |
| Age × Walkway YA Obstructed walkway | $fp_{+1}$ | $fp_{-4}$ | 0.2027 | 10052 | 48.50 | <.0001* |
| Age × Walkway YA Obstructed walkway | $fp_{+2}$ | $fp_{+3}$ | -0.0476 | 10052 | -11.41 | <.0001* |
| Age × Walkway YA Obstructed walkway | $fp_{+2}$ | $fp_{-1}$ | -0.0412 | 10052 | -9.88 | <.0001* |
| Age × Walkway YA Obstructed walkway | $fp_{+2}$ | $fp_{-2}$ | -0.0303 | 10052 | -7.27 | <.0001* |
| Age × Walkway YA Obstructed walkway | $fp_{+2}$ | $fp_{-3}$ | -0.0241 | 10052 | -5.78 | <.0001* |
| Age × Walkway YA Obstructed walkway | $fp_{+2}$ | $fp_{-4}$ | -0.0494 | 10052 | -11.84 | <.0001* |
| Age × Walkway YA Obstructed walkway | $fp_{+3}$ | $fp_{-1}$ | 0.0063 | 10052 | 1.53 | 0.13 |
| Age × Walkway YA Obstructed walkway | $fp_{+3}$ | $fp_{-2}$ | 0.0173 | 10052 | 4.14 | <.0001* |
| Age × Walkway YA Obstructed walkway | $fp_{+3}$ | $fp_{-3}$ | 0.0235 | 10052 | 5.62 | <.0001* |
| Age × Walkway YA Obstructed walkway | $fp_{+3}$ | $fp_{-4}$ | -0.0017 | 10052 | -0.43 | 0.66 |
| Age × Walkway YA Obstructed walkway | $fp_{-1}$ | $fp_{-2}$ | 0.0109 | 10052 | 2.61 | 0.01* |
| Age × Walkway YA Obstructed walkway | $fp_{-1}$ | $fp_{-3}$ | 0.0171 | 10052 | 4.10 | <.0001* |
| Age × Walkway YA Obstructed walkway | $fp_{-1}$ | $fp_{-4}$ | -0.0081 | 10052 | -1.96 | 0.05 |
| Age × Walkway YA Obstructed walkway | $fp_{-2}$ | $fp_{-3}$ | 0.0062 | 10052 | 1.49 | 0.14 |
| Age × Walkway YA Obstructed walkway | $fp_{-2}$ | $fp_{-4}$ | -0.0190 | 10052 | -4.57 | <.0001* |
| Age × Walkway YA Obstructed walkway | $fp_{-3}$ | $fp_{-4}$ | -0.0253 | 10052 | -6.05 | <.0001* |

### Appendix B

Pre-planned pairwise comparisons of simple effects following the significant age × walkway × foot placement interaction for center of mass position relative to the rear heel. Each comparison tested one independent variable while holding the other two constant. Thus, age-group comparisons were evaluated separately for each walkway and foot placement; walkway comparisons were evaluated separately for each age group and foot placement; and foot placement comparisons were evaluated separately for each age group and walkway. Least-squares mean differences are reported with estimates, degrees of freedom, t values, and p values. OA = older adults; YA = younger adults; df = degrees of freedom.

**Table 2.1. Type III Tests of Fixed Effects for center of mass position.**

| Effect | Num DF | Den DF | F Value | Pr > F |
|---|---|---|---|---|
| Age | 1 | 10052 | 513.18 | <.0001 |
| Walkway | 1 | 10052 | 73.77 | <.0001 |
| Age × Walkway | 1 | 10052 | 6.16 | 0.013 |
| fp | 6 | 10052 | 282.78 | <.0001 |
| Age × fp | 6 | 10052 | 19.75 | <.0001 |
| Walkway × fp | 6 | 10052 | 324.99 | <.0001 |
| Age × Walkway × fp | 6 | 10052 | 13.04 | <.0001 |

**Table 2.2. Pairwise comparisons between clear and obstructed walkway for least-squares center of mass position, evaluated separately for each age group and foot placement.**

| Simple Effect Level | Walkway | Walkway | Estimate | DF | t Value | Pr > \|t\| |
|---|---|---|---|---|---|---|
| OA $fp_{-4}$ | Clear Walkway | Obstructed Walkway | 0.0005 | 10052 | 0.15 | 0.88 |
| OA $fp_{-3}$ | Clear Walkway | Obstructed Walkway | 0.0124 | 10052 | 3.76 | 0.0002* |
| OA $fp_{-2}$ | Clear Walkway | Obstructed Walkway | 0.0148 | 10052 | 4.49 | <.0001* |
| OA $fp_{-1}$ | Clear Walkway | Obstructed Walkway | 0.0303 | 10052 | 9.19 | <.0001* |
| OA $fp_{+1}$ | Clear Walkway | Obstructed Walkway | 0.0896 | 10052 | 27.22 | <.0001* |
| OA $fp_{+2}$ | Clear Walkway | Obstructed Walkway | -0.0681 | 10052 | -20.69 | <.0001* |
| OA $fp_{+3}$ | Clear Walkway | Obstructed Walkway | -0.0126 | 10052 | -3.82 | 0.0001* |
| YA $fp_{-4}$ | Clear Walkway | Obstructed Walkway | 0.0040 | 10052 | 1.27 | 0.20 |
| YA $fp_{-3}$ | Clear Walkway | Obstructed Walkway | 0.0089 | 10052 | 2.83 | 0.005* |
| YA $fp_{-2}$ | Clear Walkway | Obstructed Walkway | 0.0160 | 10052 | 5.07 | <.0001* |
| YA $fp_{-1}$ | Clear Walkway | Obstructed Walkway | 0.0205 | 10052 | 6.49 | <.0001* |
| YA $fp_{+1}$ | Clear Walkway | Obstructed Walkway | 0.0518 | 10052 | 16.40 | <.0001* |
| YA $fp_{+2}$ | Clear Walkway | Obstructed Walkway | -0.0660 | 10052 | -20.91 | <.0001* |
| YA $fp_{+3}$ | Clear Walkway | Obstructed Walkway | 0.0016 | 10052 | 0.51 | 0.60 |

**Table 2.3. Pairwise comparisons between younger and older adults for least-squares center of mass position, evaluated separately for each walkway and foot placement.**

| Simple Effect Level | | Age | Age | Estimate | DF | t Value | Pr > \|t\| |
|---|---|---|---|---|---|---|---|
| Walkway × fp | Clear walkway $fp_{+1}$ | OA | YA | -0.0146 | 10052 | -4.53 | <.0001* |
| Walkway × fp | Clear walkway $fp_{+2}$ | OA | YA | -0.0258 | 10052 | -8.02 | <.0001* |
| Walkway × fp | Clear walkway $fp_{+3}$ | OA | YA | -0.0144 | 10052 | -4.48 | <.0001* |
| Walkway × fp | Clear walkway $fp_{-1}$ | OA | YA | -0.0238 | 10052 | -7.38 | <.0001* |
| Walkway × fp | Clear walkway $fp_{-2}$ | OA | YA | -0.0146 | 10052 | -4.53 | <.0001* |
| Walkway × fp | Clear walkway $fp_{-3}$ | OA | YA | -0.0185 | 10052 | -5.73 | <.0001* |
| Walkway × fp | Clear walkway $fp_{-4}$ | OA | YA | -0.0099 | 10052 | -3.07 | 0.002* |
| Walkway × fp | Obstructed Walkway $fp_{+1}$ | OA | YA | -0.0524 | 10052 | -16.24 | <.0001* |
| Walkway × fp | Obstructed Walkway $fp_{+2}$ | OA | YA | -0.0237 | 10052 | -7.37 | <.0001* |
| Walkway × fp | Obstructed Walkway $fp_{+3}$ | OA | YA | -0.0002 | 10052 | -0.07 | 0.94 |
| Walkway × fp | Obstructed Walkway $fp_{-1}$ | OA | YA | -0.0335 | 10052 | -10.41 | <.0001* |
| Walkway × fp | Obstructed Walkway $fp_{-2}$ | OA | YA | -0.0133 | 10052 | -4.15 | <.0001* |
| Walkway × fp | Obstructed Walkway $fp_{-3}$ | OA | YA | -0.0219 | 10052 | -6.80 | <.0001* |
| Walkway × fp | Obstructed Walkway $fp_{-4}$ | OA | YA | -0.0063 | 10052 | -1.97 | 0.04* |

**Table 2.4. Pairwise comparisons between foot placements for least-squares center of mass position, evaluated separately for each age group and walkway.**

| Simple Effect Level | | fp | fp | Estimate | DF | t Value | Pr > \|t\| |
|---|---|---|---|---|---|---|---|
| Age × Walkway | OA Clear walkway | $fp_{+1}$ | $fp_{+2}$ | 0.0123 | 10052 | 3.73 | 0.0002* |
| Age × Walkway | OA Clear walkway | $fp_{+1}$ | $fp_{+3}$ | 0.0230 | 10052 | 6.99 | <.0001* |
| Age × Walkway | OA Clear walkway | $fp_{+1}$ | $fp_{-1}$ | 0.0006 | 10052 | 0.21 | 0.83 |
| Age × Walkway | OA Clear walkway | $fp_{+1}$ | $fp_{-2}$ | -0.0061 | 10052 | -1.87 | 0.06 |
| Age × Walkway | OA Clear walkway | $fp_{+1}$ | $fp_{-3}$ | -0.0021 | 10052 | -0.65 | 0.51 |
| Age × Walkway | OA Clear walkway | $fp_{+1}$ | $fp_{-4}$ | -0.0005 | 10052 | -0.14 | 0.88 |
| Age × Walkway | OA Clear walkway | $fp_{+2}$ | $fp_{+3}$ | 0.0107 | 10052 | 3.25 | 0.001* |
| Age × Walkway | OA Clear walkway | $fp_{+2}$ | $fp_{-1}$ | -0.0116 | 10052 | -3.52 | 0.0004* |
| Age × Walkway | OA Clear walkway | $fp_{+2}$ | $fp_{-2}$ | -0.0184 | 10052 | -5.60 | <.0001* |
| Age × Walkway | OA Clear walkway | $fp_{+2}$ | $fp_{-3}$ | -0.0144 | 10052 | -4.38 | <.0001* |
| Age × Walkway | OA Clear walkway | $fp_{+2}$ | $fp_{-4}$ | -0.0127 | 10052 | -3.87 | 0.0001* |
| Age × Walkway | OA Clear walkway | $fp_{+3}$ | $fp_{-1}$ | -0.0223 | 10052 | -6.78 | <.0001* |
| Age × Walkway | OA Clear walkway | $fp_{+3}$ | $fp_{-2}$ | -0.0291 | 10052 | -8.85 | <.0001* |
| Age × Walkway | OA Clear walkway | $fp_{+3}$ | $fp_{-3}$ | -0.0251 | 10052 | -7.63 | <.0001* |

| | | | | | | |
|---|---|---|---|---|---|---|
| Age × Walkway OA Clear walkway | $fp_{+3}$ | $fp_{-4}$ | -0.0234 | 10052 | -7.13 | <.0001* |
| Age × Walkway OA Clear walkway | $fp_{-1}$ | $fp_{-2}$ | -0.0068 | 10052 | -2.07 | 0.04* |
| Age × Walkway OA Clear walkway | $fp_{-1}$ | $fp_{-3}$ | -0.0028 | 10052 | -0.85 | 0.39 |
| Age × Walkway OA Clear walkway | $fp_{-1}$ | $fp_{-4}$ | -0.0011 | 10052 | -0.35 | 0.72 |
| Age × Walkway OA Clear walkway | $fp_{-2}$ | $fp_{-3}$ | 0.0040 | 10052 | 1.22 | 0.22 |
| Age × Walkway OA Clear walkway | $fp_{-2}$ | $fp_{-4}$ | 0.0057 | 10052 | 1.73 | 0.08 |
| Age × Walkway OA Clear walkway | $fp_{-3}$ | $fp_{-4}$ | 0.0016 | 10052 | 0.51 | 0.61 |
| Age × Walkway OA Obstructed walkway | $fp_{+1}$ | $fp_{+2}$ | -0.1455 | 10052 | -44.18 | <.0001* |
| Age × Walkway OA Obstructed walkway | $fp_{+1}$ | $fp_{+3}$ | -0.0791 | 10052 | -24.05 | <.0001* |
| Age × Walkway OA Obstructed walkway | $fp_{+1}$ | $fp_{-1}$ | -0.0586 | 10052 | -17.82 | <.0001* |
| Age × Walkway OA Obstructed walkway | $fp_{+1}$ | $fp_{-2}$ | -0.0809 | 10052 | -24.59 | <.0001* |
| Age × Walkway OA Obstructed walkway | $fp_{+1}$ | $fp_{-3}$ | -0.0793 | 10052 | -24.10 | <.0001* |
| Age × Walkway OA Obstructed walkway | $fp_{+1}$ | $fp_{-4}$ | -0.0895 | 10052 | -27.20 | <.0001* |
| Age × Walkway OA Obstructed walkway | $fp_{+2}$ | $fp_{+3}$ | 0.0662 | 10052 | 20.12 | <.0001* |
| Age × Walkway OA Obstructed walkway | $fp_{+2}$ | $fp_{-1}$ | 0.0868 | 10052 | 26.36 | <.0001* |
| Age × Walkway OA Obstructed walkway | $fp_{+2}$ | $fp_{-2}$ | 0.0645 | 10052 | 19.59 | <.0001* |
| Age × Walkway OA Obstructed walkway | $fp_{+2}$ | $fp_{-3}$ | 0.0661 | 10052 | 20.08 | <.0001* |
| Age × Walkway OA Obstructed walkway | $fp_{+2}$ | $fp_{-4}$ | 0.0558 | 10052 | 16.97 | <.0001* |
| Age × Walkway OA Obstructed walkway | $fp_{+3}$ | $fp_{-1}$ | 0.0205 | 10052 | 6.24 | <.0001* |
| Age × Walkway OA Obstructed walkway | $fp_{+3}$ | $fp_{-2}$ | -0.0017 | 10052 | -0.54 | 0.59 |
| Age × Walkway OA Obstructed walkway | $fp_{+3}$ | $fp_{-3}$ | -0.0001 | 10052 | -0.05 | 0.96 |
| Age × Walkway OA Obstructed walkway | $fp_{+3}$ | $fp_{-4}$ | -0.0103 | 10052 | -3.15 | 0.002* |
| Age × Walkway OA Obstructed walkway | $fp_{-1}$ | $fp_{-2}$ | -0.0223 | 10052 | -6.77 | <.0001* |
| Age × Walkway OA Obstructed walkway | $fp_{-1}$ | $fp_{-3}$ | -0.0207 | 10052 | -6.29 | <.0001* |
| Age × Walkway OA Obstructed walkway | $fp_{-1}$ | $fp_{-4}$ | -0.0309 | 10052 | -9.39 | <.0001* |
| Age × Walkway OA Obstructed walkway | $fp_{-2}$ | $fp_{-3}$ | 0.0016 | 10052 | 0.49 | 0.62 |
| Age × Walkway OA Obstructed walkway | $fp_{-2}$ | $fp_{-4}$ | -0.0086 | 10052 | -2.62 | 0.009* |
| Age × Walkway OA Obstructed walkway | $fp_{-3}$ | $fp_{-4}$ | -0.0102 | 10052 | -3.10 | 0.002* |
| Age × Walkway YA Clear walkway | $fp_{+1}$ | $fp_{+2}$ | 0.0010 | 10052 | 0.33 | 0.74 |
| Age × Walkway YA Clear walkway | $fp_{+1}$ | $fp_{+3}$ | 0.0231 | 10052 | 7.34 | <.0001* |
| Age × Walkway YA Clear walkway | $fp_{+1}$ | $fp_{-1}$ | -0.0085 | 10052 | -2.70 | 0.007* |
| Age × Walkway YA Clear walkway | $fp_{+1}$ | $fp_{-2}$ | -0.0061 | 10052 | -1.95 | 0.05 |
| Age × Walkway YA Clear walkway | $fp_{+1}$ | $fp_{-3}$ | -0.0060 | 10052 | -1.91 | 0.06 |
| Age × Walkway YA Clear walkway | $fp_{+1}$ | $fp_{-4}$ | 0.0042 | 10052 | 1.34 | 0.17 |
| Age × Walkway YA Clear walkway | $fp_{+2}$ | $fp_{+3}$ | 0.0221 | 10052 | 7.01 | <.0001* |
| Age × Walkway YA Clear walkway | $fp_{+2}$ | $fp_{-1}$ | -0.0095 | 10052 | -3.03 | 0.002* |
| Age × Walkway YA Clear walkway | $fp_{+2}$ | $fp_{-2}$ | -0.0072 | 10052 | -2.28 | 0.02* |
| Age × Walkway YA Clear walkway | $fp_{+2}$ | $fp_{-3}$ | -0.0070 | 10052 | -2.23 | 0.02* |
| Age × Walkway YA Clear walkway | $fp_{+2}$ | $fp_{-4}$ | 0.0032 | 10052 | 1.02 | 0.30 |
| Age × Walkway YA Clear walkway | $fp_{+3}$ | $fp_{-1}$ | -0.0317 | 10052 | -10.03 | <.0001* |
| Age × Walkway YA Clear walkway | $fp_{+3}$ | $fp_{-2}$ | -0.0293 | 10052 | -9.29 | <.0001* |
| Age × Walkway YA Clear walkway | $fp_{+3}$ | $fp_{-3}$ | -0.0292 | 10052 | -9.24 | <.0001* |
| Age × Walkway YA Clear walkway | $fp_{+3}$ | $fp_{-4}$ | -0.0189 | 10052 | -5.99 | <.0001* |
| Age × Walkway YA Clear walkway | $fp_{-1}$ | $fp_{-2}$ | 0.0023 | 10052 | 0.75 | 0.45 |
| Age × Walkway YA Clear walkway | $fp_{-1}$ | $fp_{-3}$ | 0.0025 | 10052 | 0.79 | 0.42 |

| | | | | | | |
|---|---|---|---|---|---|---|
| Age × Walkway YA Clear walkway | $fp_{-1}$ | $fp_{-4}$ | 0.0127 | 10052 | 4.04 | <.0001* |
| Age × Walkway YA Clear walkway | $fp_{-2}$ | $fp_{-3}$ | 0.0001 | 10052 | 0.04 | 0.96 |
| Age × Walkway YA Clear walkway | $fp_{-2}$ | $fp_{-4}$ | 0.0104 | 10052 | 3.29 | 0.001* |
| Age × Walkway YA Clear walkway | $fp_{-3}$ | $fp_{-4}$ | 0.0102 | 10052 | 3.25 | 0.001* |
| Age × Walkway YA Obstructed walkway | $fp_{+1}$ | $fp_{+2}$ | -0.1168 | 10052 | -36.99 | <.0001* |
| Age × Walkway YA Obstructed walkway | $fp_{+1}$ | $fp_{+3}$ | -0.0270 | 10052 | -8.56 | <.0001* |
| Age × Walkway YA Obstructed walkway | $fp_{+1}$ | $fp_{-1}$ | -0.0398 | 10052 | -12.62 | <.0001* |
| Age × Walkway YA Obstructed walkway | $fp_{+1}$ | $fp_{-2}$ | -0.0419 | 10052 | -13.28 | <.0001* |
| Age × Walkway YA Obstructed walkway | $fp_{+1}$ | $fp_{-3}$ | -0.0488 | 10052 | -15.48 | <.0001* |
| Age × Walkway YA Obstructed walkway | $fp_{+1}$ | $fp_{-4}$ | -0.0435 | 10052 | -13.79 | <.0001* |
| Age × Walkway YA Obstructed walkway | $fp_{+2}$ | $fp_{+3}$ | 0.0898 | 10052 | 28.43 | <.0001* |
| Age × Walkway YA Obstructed walkway | $fp_{+2}$ | $fp_{-1}$ | 0.0769 | 10052 | 24.37 | <.0001* |
| Age × Walkway YA Obstructed walkway | $fp_{+2}$ | $fp_{-2}$ | 0.0748 | 10052 | 23.71 | <.0001* |
| Age × Walkway YA Obstructed walkway | $fp_{+2}$ | $fp_{-3}$ | 0.0679 | 10052 | 21.51 | <.0001* |
| Age × Walkway YA Obstructed walkway | $fp_{+2}$ | $fp_{-4}$ | 0.0732 | 10052 | 23.20 | <.0001* |
| Age × Walkway YA Obstructed walkway | $fp_{+3}$ | $fp_{-1}$ | -0.0128 | 10052 | -4.06 | <.0001* |
| Age × Walkway YA Obstructed walkway | $fp_{+3}$ | $fp_{-2}$ | -0.0149 | 10052 | -4.73 | <.0001* |
| Age × Walkway YA Obstructed walkway | $fp_{+3}$ | $fp_{-3}$ | -0.0218 | 10052 | -6.93 | <.0001* |
| Age × Walkway YA Obstructed walkway | $fp_{+3}$ | $fp_{-4}$ | -0.0165 | 10052 | -5.23 | <.0001* |
| Age × Walkway YA Obstructed walkway | $fp_{-1}$ | $fp_{-2}$ | -0.0021 | 10052 | -0.67 | 0.50 |
| Age × Walkway YA Obstructed walkway | $fp_{-1}$ | $fp_{-3}$ | -0.0090 | 10052 | -2.87 | 0.004* |
| Age × Walkway YA Obstructed walkway | $fp_{-1}$ | $fp_{-4}$ | -0.0037 | 10052 | -1.17 | 0.24 |
| Age × Walkway YA Obstructed walkway | $fp_{-2}$ | $fp_{-3}$ | -0.0069 | 10052 | -2.20 | 0.03* |
| Age × Walkway YA Obstructed walkway | $fp_{-2}$ | $fp_{-4}$ | -0.0016 | 10052 | -0.50 | 0.61 |
| Age × Walkway YA Obstructed walkway | $fp_{-3}$ | $fp_{-4}$ | 0.0053 | 10052 | 1.69 | 0.09 |

### Appendix C

Pre-planned pairwise comparisons of simple effects following the significant age × walkway × foot placement interaction for center of mass velocity. Each comparison tested one independent variable while holding the other two constant. Thus, age-group comparisons were evaluated separately for each walkway and foot placement; walkway comparisons were evaluated separately for each age group and foot placement; and foot placement comparisons were evaluated separately for each age group and walkway. Least-squares mean differences are reported with estimates, degrees of freedom, t values, and p values. OA = older adults; YA = younger adults; df = degrees of freedom.

**Table 3.1. Type III tests of fixed effects for center of mass velocity.**

| Effect | Num DF | Den DF | F Value | Pr > F |
|---|---|---|---|---|
| Age | 1 | 10052 | 359.92 | <.0001 |
| Walkway | 1 | 10052 | 589.88 | <.0001 |
| Age × Walkway | 1 | 10052 | 26.13 | <.0001 |
| fp | 6 | 10052 | 158.54 | <.0001 |
| Age × fp | 6 | 10052 | 12.02 | <.0001 |
| Walkway × fp | 6 | 10052 | 164.74 | <.0001 |
| Age × Walkway × fp | 6 | 10052 | 9.61 | <.0001 |

**Table 3.2 Pairwise comparisons between clear and obstructed walkway for least-squares for center of mass velocity, evaluated separately for each age group and foot placement.**

| Simple Effect Level | Walkway | Walkway | Estimate | DF | t Value | Pr > \|t\| |
|---|---|---|---|---|---|---|
| OA $fp_{+1}$ | Clear walkway | Obstructed walkway | 0.4399 | 10052 | 31.66 | <.0001* |
| OA $fp_{+2}$ | Clear walkway | Obstructed walkway | 0.0760 | 10052 | 5.47 | <.0001* |
| OA $fp_{+3}$ | Clear walkway | Obstructed walkway | 0.0028 | 10052 | 0.20 | 0.83 |
| OA $fp_{-1}$ | Clear walkway | Obstructed walkway | 0.1211 | 10052 | 8.72 | <.0001* |
| OA $fp_{-2}$ | Clear walkway | Obstructed walkway | 0.0700 | 10052 | 5.04 | <.0001* |
| OA $fp_{-3}$ | Clear walkway | Obstructed walkway | 0.0422 | 10052 | 3.04 | 0.002* |
| OA $fp_{-4}$ | Clear walkway | Obstructed walkway | -0.0033 | 10052 | -0.24 | 0.81 |
| YA $fp_{+1}$ | Clear walkway | Obstructed walkway | 0.2754 | 10052 | 20.66 | <.0001* |
| YA $fp_{+2}$ | Clear walkway | Obstructed walkway | 0.0373 | 10052 | 2.80 | 0.005* |
| YA $fp_{+3}$ | Clear walkway | Obstructed walkway | 0.0008 | 10052 | 0.07 | 0.94 |
| YA $fp_{-1}$ | Clear walkway | Obstructed walkway | 0.0764 | 10052 | 5.74 | <.0001* |
| YA $fp_{-2}$ | Clear walkway | Obstructed walkway | 0.0583 | 10052 | 4.38 | <.0001* |
| YA $fp_{-3}$ | Clear walkway | Obstructed walkway | 0.0290 | 10052 | 2.18 | 0.03* |
| YA $fp_{-4}$ | Clear walkway | Obstructed walkway | 0.0109 | 10052 | 0.82 | 0.41 |

**Table 3.3 Pairwise comparisons between younger and older adults for least-squares mean for center of mass velocity, evaluated separately for each walkway and foot placement.**

| Simple Effect Level | Age | Age | Estimate | DF | t Value | Pr > \|t\| |
|---|---|---|---|---|---|---|
| Walkway × fp Clear Walkway $fp_{+1}$ | OA | YA | -0.0473 | 10052 | -3.48 | 0.0005* |
| Walkway × fp Clear Walkway $fp_{+2}$ | OA | YA | -0.0680 | 10052 | -5.00 | <.0001* |
| Walkway × fp Clear Walkway $fp_{+3}$ | OA | YA | -0.0467 | 10052 | -3.44 | 0.0006* |
| Walkway × fp Clear Walkway $fp_{-1}$ | OA | YA | -0.0599 | 10052 | -4.40 | <.0001* |
| Walkway × fp Clear Walkway $fp_{-2}$ | OA | YA | -0.0441 | 10052 | -3.24 | 0.001* |
| Walkway × fp Clear Walkway $fp_{-3}$ | OA | YA | -0.0495 | 10052 | -3.64 | 0.0003* |
| Walkway × fp Clear Walkway $fp_{-4}$ | OA | YA | -0.0371 | 10052 | -2.73 | 0.006* |
| Walkway × fp Obstructed Walkway $fp_{+1}$ | OA | YA | -0.2119 | 10052 | -15.56 | <.0001* |
| Walkway × fp Obstructed Walkway $fp_{+2}$ | OA | YA | -0.1067 | 10052 | -7.84 | <.0001* |
| Walkway × fp Obstructed Walkway$fp_{+3}$ | OA | YA | -0.0487 | 10052 | -3.58 | 0.0003* |
| Walkway × fp Obstructed Walkway $fp_{-1}$ | OA | YA | -0.1046 | 10052 | -7.68 | <.0001* |
| Walkway × fp Obstructed Walkway $fp_{-2}$ | OA | YA | -0.0557 | 10052 | -4.10 | <.0001* |
| Walkway × fp Obstructed Walkway $fp_{-3}$ | OA | YA | -0.0628 | 10052 | -4.61 | <.0001* |
| Walkway × fp Obstructed Walkway $fp_{-4}$ | OA | YA | -0.0229 | 10052 | -1.69 | 0.09 |

**Table 3.4. Pairwise comparisons between foot placements for least-squares for least-squares mean for center of mass velocity, evaluated separately for each age group and walkway.**

| Simple Effect Level | fp | fp | Estimate | DF | t Value | Pr > \|t\| |
|---|---|---|---|---|---|---|
| Age × Walkway OA Clear Walkway | fp+1 | fp+2 | 0.0337 | 10052 | 2.42 | 0.01* |
| Age × Walkway OA Clear Walkway | fp+1 | fp+3 | 0.0958 | 10052 | 6.90 | <.0001* |
| Age × Walkway OA Clear Walkway | fp+1 | fp-1 | -0.0121 | 10052 | -0.88 | 0.38 |
| Age × Walkway OA Clear Walkway | fp+1 | fp-2 | -0.0214 | 10052 | -1.54 | 0.12 |
| Age × Walkway OA Clear Walkway | fp+1 | fp-3 | 0.0148 | 10052 | 1.06 | 0.28 |
| Age × Walkway OA Clear Walkway | fp+1 | fp-4 | 0.1177 | 10052 | 8.47 | <.0001* |
| Age × Walkway OA Clear Walkway | fp+2 | fp+3 | 0.0622 | 10052 | 4.48 | <.0001* |
| Age × Walkway OA Clear Walkway | fp+2 | fp-1 | -0.0458 | 10052 | -3.30 | 0.001* |
| Age × Walkway OA Clear Walkway | fp+2 | fp-2 | -0.0551 | 10052 | -3.97 | <.0001* |
| Age × Walkway OA Clear Walkway | fp+2 | fp-3 | -0.0188 | 10052 | -1.36 | 0.17 |
| Age × Walkway OA Clear Walkway | fp+2 | fp-4 | 0.0840 | 10052 | 6.04 | <.0001* |
| Age × Walkway OA Clear Walkway | fp+3 | fp-1 | -0.1080 | 10052 | -7.77 | <.0001* |
| Age × Walkway OA Clear Walkway | fp+3 | fp-2 | -0.1173 | 10052 | -8.44 | <.0001* |
| Age × Walkway OA Clear Walkway | fp+3 | fp-3 | -0.0810 | 10052 | -5.83 | <.0001* |
| Age × Walkway OA Clear Walkway | fp+3 | fp-4 | 0.0218 | 10052 | 1.57 | 0.12 |
| Age × Walkway OA Clear Walkway | fp-1 | fp-2 | -0.0092 | 10052 | -0.67 | 0.50 |
| Age × Walkway OA Clear Walkway | fp-1 | fp-3 | 0.0269 | 10052 | 1.94 | 0.05 |
| Age × Walkway OA Clear Walkway | fp-1 | fp-4 | 0.1298 | 10052 | 9.34 | <.0001* |
| Age × Walkway OA Clear Walkway | fp-2 | fp-3 | 0.0362 | 10052 | 2.61 | 0.009* |
| Age × Walkway OA Clear Walkway | fp-2 | fp-4 | 0.1391 | 10052 | 10.01 | <.0001* |
| Age × Walkway OA Clear Walkway | fp-3 | fp-4 | 0.1029 | 10052 | 7.40 | <.0001* |

| | | | | | | | |
|---|---|---|---|---|---|---|---|
| Age × Walkway | OA Obstructed Walkway | fp+1 | fp+2 | -0.3302 | 10052 | -23.76 | <.0001* |
| Age × Walkway | OA Obstructed Walkway | fp+1 | fp+3 | -0.3412 | 10052 | -24.55 | <.0001* |
| Age × Walkway | OA Obstructed Walkway | fp+1 | fp-1 | -0.3309 | 10052 | -23.82 | <.0001* |
| Age × Walkway | OA Obstructed Walkway | fp+1 | fp-2 | -0.3913 | 10052 | -28.16 | <.0001* |
| Age × Walkway | OA Obstructed Walkway | fp+1 | fp-3 | -0.3828 | 10052 | -27.55 | <.0001* |
| Age × Walkway | OA Obstructed Walkway | fp+1 | fp-4 | -0.3255 | 10052 | -23.43 | <.0001* |
| Age × Walkway | OA Obstructed Walkway | fp+2 | fp+3 | -0.0110 | 10052 | -0.79 | 0.42 |
| Age × Walkway | OA Obstructed Walkway | fp+2 | fp-1 | -0.0007 | 10052 | -0.05 | 0.95 |
| Age × Walkway | OA Obstructed Walkway | fp+2 | fp-2 | -0.0611 | 10052 | -4.40 | <.0001* |
| Age × Walkway | OA Obstructed Walkway | fp+2 | fp-3 | -0.0526 | 10052 | -3.79 | 0.0002* |
| Age × Walkway | OA Obstructed Walkway | fp+2 | fp-4 | 0.0046 | 10052 | 0.33 | 0.74 |
| Age × Walkway | OA Obstructed Walkway | fp+3 | fp-1 | 0.0102 | 10052 | 0.74 | 0.46 |
| Age × Walkway | OA Obstructed Walkway | fp+3 | fp-2 | -0.0501 | 10052 | -3.61 | 0.0003* |
| Age × Walkway | OA Obstructed Walkway | fp+3 | fp-3 | -0.0416 | 10052 | -3.00 | 0.003* |
| Age × Walkway | OA Obstructed Walkway | fp+3 | fp-4 | 0.0156 | 10052 | 1.13 | 0.26 |
| Age × Walkway | OA Obstructed Walkway | fp-1 | fp-2 | -0.0603 | 10052 | -4.34 | <.0001* |
| Age × Walkway | OA Obstructed Walkway | fp-1 | fp-3 | -0.0518 | 10052 | -3.73 | 0.0002* |
| Age × Walkway | OA Obstructed Walkway | fp-1 | fp-4 | 0.0054 | 10052 | 0.39 | 0.70 |
| Age × Walkway | OA Obstructed Walkway | fp-2 | fp-3 | 0.0084 | 10052 | 0.61 | 0.54 |
| Age × Walkway | OA Obstructed Walkway | fp-2 | fp-4 | 0.0657 | 10052 | 4.73 | <.0001* |
| Age × Walkway | OA Obstructed Walkway | fp-3 | fp-4 | 0.0572 | 10052 | 4.12 | <.0001* |
| Age × Walkway | YA Clear Walkway | fp+1 | fp+2 | 0.0130 | 10052 | 0.98 | 0.33 |
| Age × Walkway | YA Clear Walkway | fp+1 | fp+3 | 0.0964 | 10052 | 7.24 | <.0001* |
| Age × Walkway | YA Clear Walkway | fp+1 | fp-1 | -0.0247 | 10052 | -1.85 | 0.06 |
| Age × Walkway | YA Clear Walkway | fp+1 | fp-2 | -0.0182 | 10052 | -1.37 | 0.17 |
| Age × Walkway | YA Clear Walkway | fp+1 | fp-3 | 0.0126 | 10052 | 0.94 | 0.34 |
| Age × Walkway | YA Clear Walkway | fp+1 | fp-4 | 0.1279 | 10052 | 9.59 | <.0001* |
| Age × Walkway | YA Clear Walkway | fp+2 | fp+3 | 0.0834 | 10052 | 6.26 | <.0001* |
| Age × Walkway | YA Clear Walkway | fp+2 | fp-1 | -0.0377 | 10052 | -2.83 | 0.005* |
| Age × Walkway | YA Clear Walkway | fp+2 | fp-2 | -0.0312 | 10052 | -2.34 | 0.02* |
| Age × Walkway | YA Clear Walkway | fp+2 | fp-3 | -0.0004 | 10052 | -0.03 | 0.97 |
| Age × Walkway | YA Clear Walkway | fp+2 | fp-4 | 0.1149 | 10052 | 8.62 | <.0001* |
| Age × Walkway | YA Clear Walkway | fp+3 | fp-1 | -0.1212 | 10052 | -9.09 | <.0001* |
| Age × Walkway | YA Clear Walkway | fp+3 | fp-2 | -0.1147 | 10052 | -8.60 | <.0001* |
| Age × Walkway | YA Clear Walkway | fp+3 | fp-3 | -0.0838 | 10052 | -6.29 | <.0001* |
| Age × Walkway | YA Clear Walkway | fp+3 | fp-4 | 0.0314 | 10052 | 2.36 | 0.02* |
| Age × Walkway | YA Clear Walkway | fp-1 | fp-2 | 0.0064 | 10052 | 0.49 | 0.62 |
| Age × Walkway | YA Clear Walkway | fp-1 | fp-3 | 0.0373 | 10052 | 2.80 | 0.005* |
| Age × Walkway | YA Clear Walkway | fp-1 | fp-4 | 0.1526 | 10052 | 11.45 | <.0001* |
| Age × Walkway | YA Clear Walkway | fp-2 | fp-3 | 0.0308 | 10052 | 2.31 | 0.02* |
| Age × Walkway | YA Clear Walkway | fp-2 | fp-4 | 0.1461 | 10052 | 10.96 | <.0001* |
| Age × Walkway | YA Clear Walkway | fp-3 | fp-4 | 0.1153 | 10052 | 8.65 | <.0001* |
| Age × Walkway | YA Obstructed Walkway | fp+1 | fp+2 | -0.2250 | 10052 | -16.88 | <.0001* |
| Age × Walkway | YA Obstructed Walkway | fp+1 | fp+3 | -0.1780 | 10052 | -13.36 | <.0001* |
| Age × Walkway | YA Obstructed Walkway | fp+1 | fp-1 | -0.2236 | 10052 | -16.78 | <.0001* |

| | | | | | | |
|---|---|---|---|---|---|---|
| Age × Walkway YA Obstructed Walkway | fp+1 | fp-2 | -0.2352 | 10052 | -17.65 | <.0001* |
| Age × Walkway YA Obstructed Walkway | fp+1 | fp-3 | -0.2337 | 10052 | -17.54 | <.0001* |
| Age × Walkway YA Obstructed Walkway | fp+1 | fp-4 | -0.1366 | 10052 | -10.25 | <.0001* |
| Age × Walkway YA Obstructed Walkway | fp+2 | fp+3 | 0.0470 | 10052 | 3.53 | 0.0004* |
| Age × Walkway YA Obstructed Walkway | fp+2 | fp-1 | 0.0014 | 10052 | 0.11 | 0.91 |
| Age × Walkway YA Obstructed Walkway | fp+2 | fp-2 | -0.0101 | 10052 | -0.76 | 0.44 |
| Age × Walkway YA Obstructed Walkway | fp+2 | fp-3 | -0.0086 | 10052 | -0.65 | 0.51 |
| Age × Walkway YA Obstructed Walkway | fp+2 | fp-4 | 0.0884 | 10052 | 6.64 | <.0001* |
| Age × Walkway YA Obstructed Walkway | fp+3 | fp-1 | -0.0455 | 10052 | -3.42 | 0.0006* |
| Age × Walkway YA Obstructed Walkway | fp+3 | fp-2 | -0.0571 | 10052 | -4.29 | <.0001* |
| Age × Walkway YA Obstructed Walkway | fp+3 | fp-3 | -0.0557 | 10052 | -4.18 | <.0001* |
| Age × Walkway YA Obstructed Walkway | fp+3 | fp-4 | 0.0414 | 10052 | 3.11 | 0.002* |
| Age × Walkway YA Obstructed Walkway | fp-1 | fp-2 | -0.0115 | 10052 | -0.87 | 0.38 |
| Age × Walkway YA Obstructed Walkway | fp-1 | fp-3 | -0.0101 | 10052 | -0.76 | 0.45 |
| Age × Walkway YA Obstructed Walkway | fp-1 | fp-4 | 0.0870 | 10052 | 6.53 | <.0001* |
| Age × Walkway YA Obstructed Walkway | fp-2 | fp-3 | 0.0014 | 10052 | 0.11 | 0.91 |
| Age × Walkway YA Obstructed Walkway | fp-2 | fp-4 | 0.0986 | 10052 | 7.40 | <.0001* |
| Age × Walkway YA Obstructed Walkway | fp-3 | fp-4 | 0.0971 | 10052 | 7.29 | <.0001* |

### Appendix D

Pre-planned pairwise comparisons of simple effects following the significant age × walkway × step interaction for step length. Each comparison tested one independent variable while holding the other two constant. Thus, age-group comparisons were evaluated separately for each walkway and step; walkway comparisons were evaluated separately for each age group and step; and step comparisons were evaluated separately for each age group and walkway. Least-squares mean differences are reported with estimates, degrees of freedom, t values, and p values. OA = older adults; YA = younger adults; df = degrees of freedom.

**Table 4.1. Type III tests of fixed effects for step length.**

| Effect | Num DF | Den DF | F Value | Pr > F |
|---|---|---|---|---|
| Age | 1 | 8616 | 443.79 | <.0001 |
| Walkway | 1 | 8616 | 94.59 | <.0001 |
| Age × Walkway | 1 | 8616 | 5.04 | 0.0249 |
| Step | 5 | 8616 | 139.43 | <.0001 |
| Age × Step | 5 | 8616 | 8.41 | <.0001 |
| Walkway × Step | 5 | 8616 | 92.21 | <.0001 |
| Age × Walkway × Step | 5 | 8616 | 2.65 | 0.0214 |

**Table 4.2 Pairwise comparisons between clear and obstructed walkway for least-squares for step length, evaluated separately for each age group and foot placement.**

| Simple Effect Level | Walkway | Walkway | Estimate | DF | t Value | Pr > \|t\| |
|---|---|---|---|---|---|---|
| OA $Step_{+1}$ | Clear walkway | Obstructed walkway | -0.0101 | 8616 | -1.86 | 0.06 |
| OA $Step_{+2}$ | Clear walkway | Obstructed walkway | -0.0152 | 8616 | -2.80 | 0.005* |
| OA $Step_{-1}$ | Clear walkway | Obstructed walkway | 0.0049 | 8616 | 0.92 | 0.35 |
| OA $Step_{-2}$ | Clear walkway | Obstructed walkway | 0.0129 | 8616 | 2.38 | 0.01* |
| OA $Step_{-3}$ | Clear walkway | Obstructed walkway | 0.0142 | 8616 | 2.63 | 0.008* |
| OA $Step_{0}$ | Clear walkway | Obstructed walkway | -0.0757 | 8616 | -13.96 | <.0001* |
| YA $Step_{+1}$ | Clear walkway | Obstructed walkway | -0.0347 | 8616 | -6.68 | <.0001* |
| YA $Step_{+2}$ | Clear walkway | Obstructed walkway | -0.0054 | 8616 | -1.05 | 0.29 |
| YA $Step_{-1}$ | Clear walkway | Obstructed walkway | 0.0066 | 8616 | 1.29 | 0.19 |
| YA $Step_{-2}$ | Clear walkway | Obstructed walkway | 0.0081 | 8616 | 1.56 | 0.11 |
| YA $Step_{-3}$ | Clear walkway | Obstructed walkway | 0.0066 | 8616 | 1.28 | 0.20 |
| YA $Step_{0}$ | Clear walkway | Obstructed walkway | -0.0914 | 8616 | -17.57 | <.0001* |

**Table 4.3 Pairwise comparisons between younger and older adults for least-squares mean for step length, evaluated separately for each walkway and foot placement.**

| Simple Effect Level | Age | Age | Estimate | DF | t Value | Pr > \|t\| |
|---|---|---|---|---|---|---|
| Walkway × Step Clear WalkwayStep$_{+1}$ | OA | YA | -0.0394 | 8616 | -7.42 | <.0001* |
| Walkway × Step Clear Walkway Step$_{+2}$ | OA | YA | -0.0286 | 8616 | -5.39 | <.0001* |
| Walkway × Step Clear Walkway Step$_{-1}$ | OA | YA | -0.0360 | 8616 | -6.78 | <.0001* |
| Walkway × Step Clear Walkway Step$_{-2}$ | OA | YA | -0.0186 | 8616 | -3.51 | 0.0004* |
| Walkway × Step Clear Walkway Step$_{-3}$ | OA | YA | -0.0289 | 8616 | -5.45 | <.0001* |
| Walkway × Step Clear Walkway Step$_{0}$ | OA | YA | -0.0215 | 8616 | -4.05 | <.0001* |
| Walkway × Step Obstructed Walkway Step$_{+1}$ | OA | YA | -0.0640 | 8616 | -12.05 | <.0001* |
| Walkway × Step Obstructed Walkway Step$_{+2}$ | OA | YA | -0.0189 | 8616 | -3.55 | 0.0004* |
| Walkway × Step Obstructed Walkway Step$_{-1}$ | OA | YA | -0.0343 | 8616 | -6.46 | <.0001* |
| Walkway × Step Obstructed Walkway Step$_{-2}$ | OA | YA | -0.0235 | 8616 | -4.42 | <.0001* |
| Walkway × Step Obstructed Walkway Step$_{-3}$ | OA | YA | -0.0366 | 8616 | -6.89 | <.0001* |
| Walkway × Step Obstructed Walkway Step$_{0}$ | OA | YA | -0.0372 | 8616 | -7.00 | <.0001* |

**Table 4.4. Pairwise comparisons between foot placements for least-squares for least-squares mean for step length, evaluated separately for each age group and walkway.**

| Simple Effect Level | Step | Step | Estimate | DF | t Value | Pr > \|t\| |
|---|---|---|---|---|---|---|
| Age*Walkway OA Clear Walkway | Step$_{+1}$ | Step$_{+2}$ | 0.0113 | 8616 | 2.09 | 0.04* |
| Age*Walkway OA Clear Walkway | Step$_{+1}$ | Step$_{-1}$ | -0.0148 | 8616 | -2.74 | 0.006* |
| Age*Walkway OA Clear Walkway | Step$_{+1}$ | Step$_{-2}$ | -0.0286 | 8616 | -5.28 | <.0001* |
| Age*Walkway OA Clear Walkway | Step$_{+1}$ | Step$_{-3}$ | -0.0155 | 8616 | -2.86 | 0.004* |
| Age*Walkway OA Clear Walkway | Step$_{+1}$ | Step$_{0}$ | -0.0224 | 8616 | -4.14 | <.0001* |
| Age*Walkway OA Clear Walkway | Step$_{+2}$ | Step$_{-1}$ | -0.0262 | 8616 | -4.83 | <.0001* |
| Age*Walkway OA Clear Walkway | Step$_{+2}$ | Step$_{-2}$ | -0.0399 | 8616 | -7.37 | <.0001* |
| Age*Walkway OA Clear Walkway | Step$_{+2}$ | Step$_{-3}$ | -0.0268 | 8616 | -4.95 | <.0001* |
| Age*Walkway OA Clear Walkway | Step$_{+2}$ | Step$_{0}$ | -0.0338 | 8616 | -6.23 | <.0001* |
| Age*Walkway OA Clear Walkway | Step$_{-1}$ | Step$_{-2}$ | -0.0137 | 8616 | -2.53 | 0.01* |
| Age*Walkway OA Clear Walkway | Step$_{-1}$ | Step$_{-3}$ | -0.0006 | 8616 | -0.12 | 0.90 |
| Age*Walkway OA Clear Walkway | Step$_{-1}$ | Step$_{0}$ | -0.0075 | 8616 | -1.40 | 0.16 |
| Age*Walkway OA Clear Walkway | Step$_{-2}$ | Step$_{-3}$ | 0.0130 | 8616 | 2.41 | 0.01* |
| Age*Walkway OA Clear Walkway | Step$_{-2}$ | Step$_{0}$ | 0.0061 | 8616 | 1.13 | 0.25 |
| Age*Walkway OA Clear Walkway | Step$_{-3}$ | Step$_{0}$ | -0.0069 | 8616 | -1.28 | 0.20 |
| Age*Walkway OA Obstructed Walkway | Step$_{+1}$ | Step$_{+2}$ | 0.0062 | 8616 | 1.15 | 0.25 |
| Age*Walkway OA Obstructed Walkway | Step$_{+1}$ | Step$_{-1}$ | 0.0002 | 8616 | 0.04 | 0.96 |
| Age*Walkway OA Obstructed Walkway | Step$_{+1}$ | Step$_{-2}$ | -0.0056 | 8616 | -1.03 | 0.30 |
| Age*Walkway OA Obstructed Walkway | Step$_{+1}$ | Step$_{-3}$ | 0.0088 | 8616 | 1.63 | 0.10 |

| | | | | | | |
|---|---|---|---|---|---|---|
| Age*Walkway OA Obstructed Walkway | Step$_{+1}$ | Step$_{0}$ | -0.0881 | 8616 | -16.24 | <.0001* |
| Age*Walkway OA Obstructed Walkway | Step$_{+2}$ | Step$_{-1}$ | -0.0060 | 8616 | -1.11 | 0.26 |
| Age*Walkway OA Obstructed Walkway | Step$_{+2}$ | Step$_{-2}$ | -0.0118 | 8616 | -2.18 | 0.03* |
| Age*Walkway OA Obstructed Walkway | Step$_{+2}$ | Step$_{-3}$ | 0.0025 | 8616 | 0.48 | 0.63 |
| Age*Walkway OA Obstructed Walkway | Step$_{+2}$ | Step$_{0}$ | -0.0943 | 8616 | -17.39 | <.0001* |
| Age*Walkway OA Obstructed Walkway | Step$_{-1}$ | Step$_{-2}$ | -0.0058 | 8616 | -1.07 | 0.28 |
| Age*Walkway OA Obstructed Walkway | Step$_{-1}$ | Step$_{-3}$ | 0.0086 | 8616 | 1.59 | 0.11 |
| Age*Walkway OA Obstructed Walkway | Step$_{-1}$ | Step$_{0}$ | -0.0883 | 8616 | -16.28 | <.0001* |
| Age*Walkway OA Obstructed Walkway | Step$_{-2}$ | Step$_{-3}$ | 0.0144 | 8616 | 2.66 | 0.007* |
| Age*Walkway OA Obstructed Walkway | Step$_{-2}$ | Step$_{0}$ | -0.0825 | 8616 | -15.21 | <.0001* |
| Age*Walkway OA Obstructed Walkway | Step$_{-3}$ | Step$_{0}$ | -0.0969 | 8616 | -17.87 | <.0001* |
| Age*Walkway YA Clear Walkway | Step$_{+1}$ | Step$_{+2}$ | 0.0221 | 8616 | 4.25 | <.0001* |
| Age*Walkway YA Clear Walkway | Step$_{+1}$ | Step$_{-1}$ | -0.0115 | 8616 | -2.21 | 0.03* |
| Age*Walkway YA Clear Walkway | Step$_{+1}$ | Step$_{-2}$ | -0.0078 | 8616 | -1.51 | 0.13 |
| Age*Walkway YA Clear Walkway | Step$_{+1}$ | Step$_{-3}$ | -0.0050 | 8616 | -0.98 | 0.32 |
| Age*Walkway YA Clear Walkway | Step$_{+1}$ | Step$_{0}$ | -0.0045 | 8616 | -0.88 | 0.38 |
| Age*Walkway YA Clear Walkway | Step$_{+2}$ | Step$_{-1}$ | -0.0336 | 8616 | -6.46 | <.0001* |
| Age*Walkway YA Clear Walkway | Step$_{+2}$ | Step$_{-2}$ | -0.0300 | 8616 | -5.76 | <.0001* |
| Age*Walkway YA Clear Walkway | Step$_{+2}$ | Step$_{-3}$ | -0.0272 | 8616 | -5.23 | <.0001* |
| Age*Walkway YA Clear Walkway | Step$_{+2}$ | Step$_{0}$ | -0.0266 | 8616 | -5.13 | <.0001* |
| Age*Walkway YA Clear Walkway | Step$_{-1}$ | Step$_{-2}$ | 0.0036 | 8616 | 0.70 | 0.48 |
| Age*Walkway YA Clear Walkway | Step$_{-1}$ | Step$_{-3}$ | 0.0064 | 8616 | 1.23 | 0.21 |
| Age*Walkway YA Clear Walkway | Step$_{-1}$ | Step$_{0}$ | 0.0069 | 8616 | 1.33 | 0.18 |
| Age*Walkway YA Clear Walkway | Step$_{-2}$ | Step$_{-3}$ | 0.0027 | 8616 | 0.53 | 0.59 |
| Age*Walkway YA Clear Walkway | Step$_{-2}$ | Step$_{0}$ | 0.0033 | 8616 | 0.64 | 0.52 |
| Age*Walkway YA Clear Walkway | Step$_{-3}$ | Step$_{0}$ | 0.0005 | 8616 | 0.10 | 0.91 |
| Age*Walkway YA Obstructed Walkway | Step$_{+1}$ | Step$_{+2}$ | 0.0514 | 8616 | 9.88 | <.0001* |
| Age*Walkway YA Obstructed Walkway | Step$_{+1}$ | Step$_{-1}$ | 0.0299 | 8616 | 5.75 | <.0001* |
| Age*Walkway YA Obstructed Walkway | Step$_{+1}$ | Step$_{-2}$ | 0.0349 | 8616 | 6.72 | <.0001* |
| Age*Walkway YA Obstructed Walkway | Step$_{+1}$ | Step$_{-3}$ | 0.0363 | 8616 | 6.98 | <.0001* |
| Age*Walkway YA Obstructed Walkway | Step$_{+1}$ | Step$_{0}$ | -0.0612 | 8616 | -11.77 | <.0001* |
| Age*Walkway YA Obstructed Walkway | Step$_{+2}$ | Step$_{-1}$ | -0.0214 | 8616 | -4.13 | <.0001* |
| Age*Walkway YA Obstructed Walkway | Step$_{+2}$ | Step$_{-2}$ | -0.0164 | 8616 | -3.16 | 0.002* |
| Age*Walkway YA Obstructed Walkway | Step$_{+2}$ | Step$_{-3}$ | -0.0151 | 8616 | -2.90 | 0.004* |
| Age*Walkway YA Obstructed Walkway | Step$_{+2}$ | Step$_{0}$ | -0.112 | 8616 | -21.65 | <.0001* |
| Age*Walkway YA Obstructed Walkway | Step$_{-1}$ | Step$_{-2}$ | 0.0050 | 8616 | 0.97 | 0.33 |
| Age*Walkway YA Obstructed Walkway | Step$_{-1}$ | Step$_{-3}$ | 0.0063 | 8616 | 1.22 | 0.22 |
| Age*Walkway YA Obstructed Walkway | Step$_{-1}$ | Step$_{0}$ | -0.0911 | 8616 | -17.52 | <.0001* |
| Age*Walkway YA Obstructed Walkway | Step$_{-2}$ | Step$_{-3}$ | 0.0013 | 8616 | 0.25 | 0.80 |
| Age*Walkway YA Obstructed Walkway | Step$_{-2}$ | Step$_{0}$ | -0.0962 | 8616 | -18.49 | <.0001* |
| Age*Walkway YA Obstructed Walkway | Step$_{-3}$ | Step$_{0}$ | -0.0975 | 8616 | -18.74 | <.0001* |

### Appendix E

Pre-planned pairwise comparisons of simple effects following the significant walkway × step interaction for the synergy index. Each comparison tested one independent variable while holding the other constant. Thus, walkway comparisons were evaluated separately at each step, and step comparisons were evaluated separately for each walkway. Least-squares mean differences are reported with estimates, degrees of freedom, t values, and p values. df = degrees of freedom.

**Table 5.1 Type III tests of fixed effects for the synergy index.**

| Effect | Num DF | Den DF | F Value | Pr > F |
|---|---|---|---|---|
| Age | 1 | 46 | 0.02 | 0.88 |
| Walkway | 1 | 506 | 9.64 | 0.002* |
| Age × Walkway | 1 | 506 | 0.12 | 0.72 |
| Step | 5 | 506 | 8.29 | <.0001* |
| Age × Step | 5 | 506 | 0.45 | 0.81 |
| Walkway × Step | 5 | 506 | 3.75 | 0.002* |
| Age × Walkway × Step | 5 | 506 | 0.75 | 0.58 |

**Table 5.2. Pairwise comparisons between clear and obstructed walkway for least-squares mean synergy index values, evaluated separately for each step.**

| Simple Effect Level | Walkway | Walkway | Estimate | DF | t Value | Pr > \|t\| |
|---|---|---|---|---|---|---|
| Step $Step_{+1}$ | Clear Walkway | Obstructed Walkway | 0.1801 | 506 | 2.95 | 0.003* |
| Step $Step_{+2}$ | Clear Walkway | Obstructed Walkway | -0.0193 | 506 | -0.32 | 0.75 |
| Step $Step_{-1}$ | Clear Walkway | Obstructed Walkway | -0.0353 | 506 | -0.58 | 0.56 |
| Step $Step_{-2}$ | Clear Walkway | Obstructed Walkway | 0.0706 | 506 | 1.16 | 0.24 |
| Step $Step_{-3}$ | Clear Walkway | Obstructed Walkway | 0.0098 | 506 | 0.16 | 0.87 |
| Step $Step_{0}$ | Clear Walkway | Obstructed Walkway | 0.2581 | 506 | 4.23 | <.0001* |

**Table 5.3. Pairwise comparisons between steps for least-squares mean synergy index values, evaluated separately for each walkway.**

| Simple Effect Level | Step | Step | Estimate | DF | t Value | Pr > \|t\| |
|---|---|---|---|---|---|---|
| Walkway Clear Walkway | $Step_{+1}$ | $Step_{+2}$ | -0.0305 | 506 | -0.50 | 0.61 |
| Walkway Clear Walkway | $Step_{+1}$ | $Step_{-1}$ | 0.0863 | 506 | 1.42 | 0.15 |
| Walkway Clear Walkway | $Step_{+1}$ | $Step_{-2}$ | 0.1016 | 506 | 1.67 | 0.09 |
| Walkway Clear Walkway | $Step_{+1}$ | $Step_{-3}$ | 0.0660 | 506 | 1.08 | 0.27 |
| Walkway Clear Walkway | $Step_{+1}$ | $Step_{0}$ | 0.0974 | 506 | 1.60 | 0.11 |
| Walkway Clear Walkway | $Step_{+2}$ | $Step_{-1}$ | 0.1169 | 506 | 1.92 | 0.06 |
| Walkway Clear Walkway | $Step_{+2}$ | $Step_{-2}$ | 0.1322 | 506 | 2.17 | 0.03* |
| Walkway Clear Walkway | $Step_{+2}$ | $Step_{-3}$ | 0.0966 | 506 | 1.58 | 0.11 |
| Walkway Clear Walkway | $Step_{+2}$ | $Step_{0}$ | 0.1281 | 506 | 2.10 | 0.04* |

| | | | | | | |
|---|---|---|---|---|---|---|
| Walkway Clear Walkway | Step$_{-1}$ | Step$_{-2}$ | 0.0152 | 506 | 0.25 | 0.80 |
| Walkway Clear Walkway | Step$_{-1}$ | Step$_{-3}$ | -0.0203 | 506 | -0.33 | 0.73 |
| Walkway Clear Walkway | Step$_{-1}$ | Step$_{0}$ | 0.0111 | 506 | 0.18 | 0.85 |
| Walkway Clear Walkway | Step$_{-2}$ | Step$_{-3}$ | -0.0355 | 506 | -0.58 | 0.56 |
| Walkway Clear Walkway | Step$_{-2}$ | Step$_{0}$ | -0.0041 | 506 | -0.07 | 0.94 |
| Walkway Clear Walkway | Step$_{-3}$ | Step$_{0}$ | 0.0314 | 506 | 0.51 | 0.60 |
| Walkway Obstructed Walkway | Step$_{+1}$ | Step$_{+2}$ | -0.2301 | 506 | -3.77 | 0.0002* |
| Walkway Obstructed Walkway | Step$_{+1}$ | Step$_{-1}$ | -0.1291 | 506 | -2.12 | 0.03* |
| Walkway Obstructed Walkway | Step$_{+1}$ | Step$_{-2}$ | -0.0078 | 506 | -0.13 | 0.89 |
| Walkway Obstructed Walkway | Step$_{+1}$ | Step$_{-3}$ | -0.1042 | 506 | -1.71 | 0.08 |
| Walkway Obstructed Walkway | Step$_{+1}$ | Step$_{0}$ | 0.1754 | 506 | 2.87 | 0.004* |
| Walkway Obstructed Walkway | Step$_{+2}$ | Step$_{-1}$ | 0.1010 | 506 | 1.65 | 0.09 |
| Walkway Obstructed Walkway | Step$_{+2}$ | Step$_{-2}$ | 0.2222 | 506 | 3.64 | 0.0003* |
| Walkway Obstructed Walkway | Step$_{+2}$ | Step$_{-3}$ | 0.1259 | 506 | 2.06 | 0.04* |
| Walkway Obstructed Walkway | Step$_{+2}$ | Step$_{0}$ | 0.4055 | 506 | 6.65 | <.0001* |
| Walkway Obstructed Walkway | Step$_{-1}$ | Step$_{-2}$ | 0.1212 | 506 | 1.99 | 0.05 |
| Walkway Obstructed Walkway | Step$_{-1}$ | Step$_{-3}$ | 0.0249 | 506 | 0.41 | 0.68 |
| Walkway Obstructed Walkway | Step$_{-1}$ | Step$_{0}$ | 0.3045 | 506 | 4.99 | <.0001* |
| Walkway Obstructed Walkway | Step$_{-2}$ | Step$_{-3}$ | -0.0963 | 506 | -1.58 | 0.11 |
| Walkway Obstructed Walkway | Step$_{-2}$ | Step$_{0}$ | 0.1833 | 506 | 3.00 | 0.003* |
| Walkway Obstructed Walkway | Step$_{-3}$ | Step$_{0}$ | 0.2796 | 506 | 4.58 | <.0001* |

**Appendix F**

Pre-planned pairwise comparisons of simple effects following the significant walkway × step interaction for the $V_{UCM}$. Each comparison tested one independent variable while holding the other constant. Thus, walkway comparisons were evaluated separately at each step, and step comparisons were evaluated separately for each walkway. Least-squares mean differences are reported with estimates, degrees of freedom, t values, and p values. df = degrees of freedom.

**Table 6.1 Type III tests of fixed effects for the $V_{UCM}$.**

| Effect | Num DF | Den DF | F Value | Pr > F |
|---|---|---|---|---|
| Age | 1 | 46 | 1.21 | 0.27 |
| Walkway | 1 | 506 | 46.66 | <.0001* |
| Age × Walkway | 1 | 506 | 0.01 | 0.92 |
| Step | 5 | 506 | 6.48 | <.0001* |
| Age × Step | 5 | 506 | 0.21 | 0.95 |
| Walkway ×Step | 5 | 506 | 8.77 | <.0001* |
| Age × Walkway × Step | 5 | 506 | 1.27 | 0.27 |

**Table 6.2. Pairwise comparisons between clear and obstructed walkway for least-squares $V_{UCM}$ values, evaluated separately for each step.**

| Simple Effect Level | Walkway | Walkway | Estimate | DF | t Value | Pr > \|t\| |
|---|---|---|---|---|---|---|
| Step $Step_{-3}$ | Clear Walkway | Obstructed Walkway | 0.000088 | 506 | 0.46 | 0.64 |
| Step $Step_{-2}$ | Clear Walkway | Obstructed Walkway | -0.00002 | 506 | -0.10 | 0.92 |
| Step $Step_{-1}$ | Clear Walkway | Obstructed Walkway | -0.00037 | 506 | -1.93 | 0.05 |
| Step $Step_{0}$ | Clear Walkway | Obstructed Walkway | -0.00121 | 506 | -6.32 | <.0001* |
| Step $Step_{+1}$ | Clear Walkway | Obstructed Walkway | -0.00122 | 506 | -6.36 | <.0001* |
| Step $Step_{+2}$ | Clear Walkway | Obstructed Walkway | -0.00048 | 506 | -2.49 | 0.01* |

**Table 6.3. Pairwise comparisons between steps for least-squares $V_{UCM}$ values, evaluated separately for each walkway.**

| Simple Effect Level | Step | Step | Estimate | DF | t Value | Pr > \|t\| |
|---|---|---|---|---|---|---|
| Walkway Clear Walkway | $Step_{+1}$ | $Step_{+2}$ | -0.0004 | 506 | -2.25 | 0.02* |
| Walkway Clear Walkway | $Step_{+1}$ | $Step_{-1}$ | -0.0001 | 506 | -0.80 | 0.42 |
| Walkway Clear Walkway | $Step_{+1}$ | $Step_{-2}$ | -0.0001 | 506 | -0.71 | 0.48 |
| Walkway Clear Walkway | $Step_{+1}$ | $Step_{-3}$ | -0.0002 | 506 | -1.22 | 0.22 |
| Walkway Clear Walkway | $Step_{+1}$ | $Step_{0}$ | -0.0008 | 506 | -0.36 | 0.71 |
| Walkway Clear Walkway | $Step_{+2}$ | $Step_{-1}$ | 0.0002 | 506 | 1.46 | 0.14 |
| Walkway Clear Walkway | $Step_{+2}$ | $Step_{-2}$ | 0.0002 | 506 | 1.55 | 0.12 |
| Walkway Clear Walkway | $Step_{+2}$ | $Step_{-3}$ | 0.0001 | 506 | 1.03 | 0.30 |
| Walkway Clear Walkway | $Step_{+2}$ | $Step_{0}$ | 0.0003 | 506 | 1.89 | 0.06 |

| | | | | | | |
|---|---|---|---|---|---|---|
| Walkway Clear Walkway | $Step_{-1}$ | $Step_{-2}$ | 0.00001 | 506 | 0.09 | 0.92 |
| Walkway Clear Walkway | $Step_{-1}$ | $Step_{-3}$ | -0.00008 | 506 | -0.43 | 0.67 |
| Walkway Clear Walkway | $Step_{-1}$ | $Step_{0}$ | 0.00008 | 506 | 0.43 | 0.66 |
| Walkway Clear Walkway | $Step_{-2}$ | $Step_{-3}$ | -0.0001 | 506 | -0.52 | 0.60 |
| Walkway Clear Walkway | $Step_{-2}$ | $Step_{0}$ | 0.00006 | 506 | 0.34 | 0.73 |
| Walkway Clear Walkway | $Step_{-3}$ | $Step_{0}$ | 0.0001 | 506 | 0.86 | 0.39 |
| Walkway Obstructed Walkway | $Step_{+1}$ | $Step_{+2}$ | 0.0003 | 506 | 1.62 | 0.10 |
| Walkway Obstructed Walkway | $Step_{+1}$ | $Step_{-1}$ | 0.0006 | 506 | 3.63 | 0.0003* |
| Walkway Obstructed Walkway | $Step_{+1}$ | $Step_{-2}$ | 0.0010 | 506 | 5.56 | <.0001* |
| Walkway Obstructed Walkway | $Step_{+1}$ | $Step_{-3}$ | 0.0010 | 506 | 5.59 | <.0001* |
| Walkway Obstructed Walkway | $Step_{+1}$ | $Step_{0}$ | -0.00006 | 506 | -0.33 | 0.74 |
| Walkway Obstructed Walkway | $Step_{+2}$ | $Step_{-1}$ | 0.0003 | 506 | 2.02 | 0.04* |
| Walkway Obstructed Walkway | $Step_{+2}$ | $Step_{-2}$ | 0.0007 | 506 | 3.94 | <.0001* |
| Walkway Obstructed Walkway | $Step_{+2}$ | $Step_{-3}$ | 0.0007 | 506 | 3.98 | <.0001* |
| Walkway Obstructed Walkway | $Step_{+2}$ | $Step_{0}$ | -0.0003 | 506 | -1.94 | 0.05 |
| Walkway Obstructed Walkway | $Step_{-1}$ | $Step_{-2}$ | 0.0003 | 506 | 1.92 | 0.05 |
| Walkway Obstructed Walkway | $Step_{-1}$ | $Step_{-3}$ | 0.0003 | 506 | 1.96 | 0.05 |
| Walkway Obstructed Walkway | $Step_{-1}$ | $Step_{0}$ | -0.0007 | 506 | -3.96 | <.0001* |
| Walkway Obstructed Walkway | $Step_{-2}$ | $Step_{-3}$ | 7.17E-6 | 506 | 0.04 | 0.97 |
| Walkway Obstructed Walkway | $Step_{-2}$ | $Step_{0}$ | -0.0011 | 506 | -5.88 | <.0001* |
| Walkway Obstructed Walkway | $Step_{-3}$ | $Step_{0}$ | -0.0011 | 506 | -5.92 | <.0001* |

### Appendix G

Pre-planned pairwise comparisons of simple effects following the significant walkway × step interaction for the $V_{ORT}$. Each comparison tested one independent variable while holding the other constant. Thus, walkway comparisons were evaluated separately at each step, and step comparisons were evaluated separately for each walkway. Least-squares mean differences are reported with estimates, degrees of freedom, t values, and p values. df = degrees of freedom.

**Table 7.1 Type III tests of fixed effects for the $V_{ORT}$.**

| Effect | Num DF | Den DF | F Value | Pr > F |
|---|---|---|---|---|
| Age | 1 | 45.51 | 1.65 | 0.20 |
| Walkway | 1 | 517.6 | 50.14 | <.0001* |
| Age × Walkway | 1 | 517.6 | 3.11 | 0.07 |
| Step | 5 | 517.6 | 10.94 | <.0001* |
| Age × Step | 5 | 517.6 | 1.16 | 0.32 |
| Walkway × Step | 5 | 517.6 | 12.59 | <.0001* |
| Age × Walkway × Step | 5 | 517.6 | 1.00 | 0.41 |

**Table 7.2. Pairwise comparisons between clear and obstructed walkway for least-squares $V_{ORT}$ values, evaluated separately for each step.**

| Simple Effect Level | Walkway | Walkway | Estimate | DF | t Value | Pr > \|t\| |
|---|---|---|---|---|---|---|
| Step $step_{+1}$ | Clear Walkway | Obstructed Walkway | -0.00026 | 517.6 | -4.10 | <.0001* |
| Step $step_{+2}$ | Clear Walkway | Obstructed Walkway | -0.00011 | 517.6 | -1.69 | 0.09 |
| Step $step_{-1}$ | Clear Walkway | Obstructed Walkway | -0.00006 | 517.6 | -0.89 | 0.37 |
| Step $step_{-2}$ | Clear Walkway | Obstructed Walkway | -0.00007 | 517.6 | -1.09 | 0.27 |
| Step $step_{-3}$ | Clear Walkway | Obstructed Walkway | -1.04E-6 | 517.6 | -0.02 | 0.98 |
| Step $step_{0}$ | Clear Walkway | Obstructed Walkway | -0.00062 | 517.6 | -9.57 | <.0001* |

**Table 7.3. Pairwise comparisons between steps for least-squares $V_{ORT}$ values, evaluated separately for each walkway.**

| Simple Effect Level | Step | Step | Estimate | DF | t Value | Pr > \|t\| |
|---|---|---|---|---|---|---|
| Walkway Clear Walkway | $step_{+1}$ | $step_{+2}$ | -0.00002 | 517.6 | -0.38 | 0.70 |
| Walkway Clear Walkway | $step_{+1}$ | $step_{-1}$ | -0.00002 | 517.6 | -0.31 | 0.75 |
| Walkway Clear Walkway | $step_{+1}$ | $step_{-2}$ | -0.00002 | 517.6 | -0.30 | 0.76 |
| Walkway Clear Walkway | $step_{+1}$ | $step_{-3}$ | -0.00003 | 517.6 | -0.48 | 0.63 |
| Walkway Clear Walkway | $step_{+1}$ | $step_{0}$ | -7.22E-6 | 517.6 | -0.11 | 0.91 |
| Walkway Clear Walkway | $step_{+2}$ | $step_{-1}$ | 4.414E-6 | 517.6 | 0.07 | 0.94 |
| Walkway Clear Walkway | $step_{+2}$ | $step_{-2}$ | 4.966E-6 | 517.6 | 0.08 | 0.93 |
| Walkway Clear Walkway | $step_{+2}$ | $step_{-3}$ | -6.58E-6 | 517.6 | -0.10 | 0.91 |
| Walkway Clear Walkway | $step_{+2}$ | $step_{0}$ | 0.000017 | 517.6 | 0.26 | 0.79 |
| Walkway Clear Walkway | $step_{-1}$ | $step_{-2}$ | 5.528E-7 | 517.6 | 0.01 | 0.99 |

| | | | | | | |
|---|---|---|---|---|---|---|
| Walkway Clear Walkway | $step_{-1}$ | $step_{-3}$ | -0.00001 | 517.6 | -0.17 | 0.86 |
| Walkway Clear Walkway | $step_{-1}$ | $step_{0}$ | 0.000013 | 517.6 | 0.19 | 0.84 |
| Walkway Clear Walkway | $step_{-2}$ | $step_{-3}$ | -0.00001 | 517.6 | -0.18 | 0.85 |
| Walkway Clear Walkway | $step_{-2}$ | $step_{0}$ | 0.000012 | 517.6 | 0.19 | 0.85 |
| Walkway Clear Walkway | $step_{-3}$ | $step_{0}$ | 0.000024 | 517.6 | 0.37 | 0.71 |
| Walkway Obstructed Walkway | $step_{+1}$ | $step_{+2}$ | 0.000131 | 517.6 | 2.03 | 0.04* |
| Walkway Obstructed Walkway | $step_{+1}$ | $step_{-1}$ | 0.000188 | 517.6 | 2.90 | 0.004* |
| Walkway Obstructed Walkway | $step_{+1}$ | $step_{-2}$ | 0.000175 | 517.6 | 2.70 | 0.007* |
| Walkway Obstructed Walkway | $step_{+1}$ | $step_{-3}$ | 0.000233 | 517.6 | 3.60 | 0.0003* |
| Walkway Obstructed Walkway | $step_{+1}$ | $step_{0}$ | -0.00036 | 517.6 | -5.58 | <.0001* |
| Walkway Obstructed Walkway | $step_{+2}$ | $step_{-1}$ | 0.000056 | 517.6 | 0.87 | 0.38 |
| Walkway Obstructed Walkway | $step_{+2}$ | $step_{-2}$ | 0.000043 | 517.6 | 0.67 | 0.50 |
| Walkway Obstructed Walkway | $step_{+2}$ | $step_{-3}$ | 0.000101 | 517.6 | 1.57 | 0.11 |
| Walkway Obstructed Walkway | $step_{+2}$ | $step_{0}$ | -0.00049 | 517.6 | -7.62 | <.0001* |
| Walkway Obstructed Walkway | $step_{-1}$ | $step_{-2}$ | -0.00001 | 517.6 | -0.20 | 0.84 |
| Walkway Obstructed Walkway | $step_{-1}$ | $step_{-3}$ | 0.000045 | 517.6 | 0.70 | 0.48 |
| Walkway Obstructed Walkway | $step_{-1}$ | $step_{0}$ | -0.00055 | 517.6 | -8.48 | <.0001* |
| Walkway Obstructed Walkway | $step_{-2}$ | $step_{-3}$ | 0.000058 | 517.6 | 0.90 | 0.36 |
| Walkway Obstructed Walkway | $step_{-2}$ | $step_{0}$ | -0.00054 | 517.6 | -8.28 | <.0001* |
| Walkway Obstructed Walkway | $step_{-3}$ | $step_{0}$ | -0.00059 | 517.6 | -9.18 | <.0001* |
